# Intramembrane protease SPPL2b deficiency linked to synaptic proteostasis remodeling with brain aging

**DOI:** 10.64898/2026.07.27.740894

**Authors:** Cristina Contini, Alessandra Schirru, Greca Lai, Giorgia Zodio, Jack Badman, Bjorn R.V. Bakker, Ylenia Lai, Per Nilsson, Giacomo Diaz, Tiziana Cabras, Simone Tambaro

## Abstract

Signal peptide peptidase-like 2b (SPPL2b) is a brain-enriched intramembrane protease implicated in synaptic function, immune signaling, and neuronal development, but its physiological role in brain and neuronal homeostasis remains poorly understood. We investigated the impact of constitutive *Sppl2b* deletion on the brain proteome and neuronal function using quantitative shotgun proteomics in cortex and hippocampus from wild-type and SPPL2b-deficient mice at 3 and 12 months, combined with western blot, immunofluorescence, dendritic spine analysis, and behavioral testing. SPPL2b deficiency was associated with alterations in biological processes linked to axonal transport and synaptic vesicle trafficking, including consistent upregulation of the kinesin motor protein KIF1A and vesicle-associated membrane protein 2 (VAMP2). SPPL2b deletion also attenuated age-associated proteomic changes in synaptic pathways and was accompanied by increased dendritic spine density, morphological remodeling of pyramidal neurons, increased locomotor activity, and reduced anxiety-like behavior. These findings reveal a previously unrecognized association between SPPL2b and synaptic proteome remodeling during aging and nominate SPPL2b-dependent pathways as candidates for further mechanistic and functional investigation in neurodegenerative disease.

## Background

Signal peptide peptidase (SPP) and signal peptide peptidase–like (SPPLs) proteins are integral membrane aspartyl proteases belonging to the GxGD–aspartyl intramembrane protease family (Weihofen *et al*, 2002; Wolfe *et al*, 1999). SPP/SPPLs were discovered by sequence homology analysis due to their shared homology with the presenilin 1 and 2 proteins (PSEN1 and PSEN2)(Ponting *et al*, 2002; Weihofen *et al*, 2002). PSEN1 and PSEN2 are the catalytic components of the γ–secretase complex(De Strooper *et al*, 2012), and cleave type I transmembrane proteins, including the amyloid-beta precursor protein (APP), which plays a crucial role in the onset of Alzheimer’s disease (AD)(Braggin *et al*, 2019; Kelleher & Shen, 2017). Beyond their role in APP processing, presenilins have also been shown to be implicated in Notch signaling, which regulates gene transcription and controls cell fate decisions and development(Ray *et al*, 1999). Overall, more than one hundred type I intramembrane proteins have so far been identified as substrates processed by presenilins(De Strooper *et al*, 1999; Haapasalo & Kovacs, 2011). On the other hand, SPP and SPPL proteins, despite being evolutionarily conserved and widely expressed, remain far less understood. The SPP/SPPLs family consists of five members: SPP, SPPL2a, SPPL2b, SPPL2c, and SPPL3(Mentrup *et al*, 2020). These enzymes share structural similarities with presenilins but differ significantly in their orientation, localization, substrate specificity, and functions(Mentrup *et al*, 2017). Unlike γ–secretase, which forms a large multiprotein complex, SPP and SPPL proteases operate as stand–alone enzymes and cleave type II transmembrane substrates, releasing an intracellular peptide, a process essential for cellular signaling and homeostasis(Badman *et al*, 2025). Among them, SPPL2b has emerged as particularly intriguing due to its predominant expression in the brain and its putative links to neurodegenerative processes(Del Campo *et al*, 2014; Fluhrer *et al*, 2006; Schneppenheim *et al*, 2014). SPPL2b and its close homolog SPPL2a cleaves several substrates, including integral membrane protein 2B (ITM2B or BRI2), cluster of differentiation 74 (CD74), tumor necrosis factor alpha (TNFα), dectin–1 (also known as C-type lectin domain family 7 member A, CLEC7A), lectin-like oxidized low-density lipoprotein receptor-1 (LOX–1), transmembrane protein 106B (TMEM106B), neuregulin–1, and Vesicle–associated membrane protein 1/2 (VAMP1, VAMP2)(Ballin *et al*, 2023). Notably, transferrin receptor 1 (Tfr–1) is cleaved specifically by SPPL2b(Zahn *et al*, 2013). SPPL2b is mainly localized to the plasma membrane and is highly expressed in neurons and glia cells, in contrast to its homolog SPPL2a, which is present in endosomal compartments of peripheral tissues such as the immune system tissues, myeloid and lymphoid cells, liver, kidney, and lung(Schneppenheim *et al*, 2014; Behnke *et al*, 2011; Friedmann *et al*, 2006). In the brain, SPPL2b substrates are associated with distinct functions, such as 1) amyloid production (BRI2, CD74), 2) synaptic function (NRG1, VAMP1, and VAMP2), and 3) neuroinflammation (CD74, LOX1, TNFα, and Dectin–1)(Ballin *et al*, 2023). Recent studies have highlighted SPPL2b’s involvement in APP processing(Maccioni *et al*, 2024) that the genetic deletion of SPPL2b alters APP cleavage and markedly reduces amyloid–β (Aβ) generation(Maccioni *et al*, 2024). This effect may be mediated through SPPL2b’s regulation of BRI2, which functions as a neuronal membrane protein that modulates APP processing(Matsuda *et al*, 2008; Fotinopoulou *et al*, 2005). SPPL2b deficiency increases BRI2 accumulation, thereby protecting APP from cleavage and limiting Aβ production. Notably, SPPL2b gene and protein expression levels have been shown to be altered during Aβ and Tau pathology in mouse models(Zhuang *et al*, 2019) and in human AD brains(Del Campo *et al*, 2014). Overall, SPPL2b’s involvement in neuronal homeostasis and synaptic maintenance highlights its broader role in brain health, extending its functional properties beyond amyloid processing(Badman *et al*, 2025). Genome–wide association studies have indeed linked SPPL2b expression to alcohol dependence and altered SPPL2b signaling has been implicated in Parkinson’s disease and hypertension(Bost *et al*, 2022; Nalls *et al*, 2019). These findings underscore its broader physiological and pathological relevance and highlight SPPL2b important role in brain neuronal processes and inflammatory pathways.

To elucidate the functional networks regulated by SPPL2b, we applied here an untargeted shotgun proteomic approach using high–resolution mass spectrometry (HR–MS) and label–free quantification. Given that SPPL2b impacts multiple substrates, involved in neuronal communication and immune signaling, we hypothesized that its deletion would trigger broad changes in protein expression, regulation and clearance affecting synaptic organization, cytoskeletal stability, and inflammatory regulation. To capture early and age–dependent effects of SPPL2b loss, and to highlight quali–/quantitative differences at proteomic level, we profiled the cortical and hippocampal proteomes of SPPL2b knockout (KO) and wild–type (WT) mice at 3 and 12-months of age.

Moreover, functional analysis of proteomic data was utilized to identify the most affected biological processes by SPPL2b delation. Among the most significant results, we focused on validating proteins of interest using western blot and immunofluorescence and we assessed dendritic spine density and neuronal morphology by Golgi staining analysis. Finally, behavioral assays were performed to evaluate impairments of locomotion and anxiolytic–like behavior induced by *Sppl2b* deletion. Our findings shed light on the multifaceted roles of SPPL2b in brain physiology and its role in modulating axonal transport and synaptic plasticity.

## Results

### Shotgun Proteomic Investigation

The explorative proteomic approach applied in this study identified 2129 proteins in the cortex and 2642 proteins in the hippocampus from WT and *Sppl2b–*KO mice. To ensure high degree of confidence in the statistical comparisons related to both aging (3– and 12–month–old mice) and genotype (WT and *Sppl2b –*KO) conditions, a filtering process with stringent criteria was applied on the proteomic data as shown in Fig. 1A–C (for full detail see the method section: “Statistic and data Filtering”). This filtering process led to the selection of 847 proteins in the cortex, and 490 in the hippocampus, each showing significant alterations related to age and/or the *Sppl2b* genotype (Fig. 1B, Source Data SD1–SD5). Selected proteins were categorized based on different affecting conditions: (1) “A” proteins, affected only by age (significant differences between 3– and 12–month– old mice); (2) “S” proteins, affected only by the deletion of the *Sppl2b* gene (significant difference between KO and WT); (3) “AS” proteins, affected by both age and gene and with significant changes in relation to both factors. These three protein categories were further divided in subcategories depending on whether protein levels increased or decreased (Fig. 1C, Source Data SD1–SD5).

**Figure 1.**
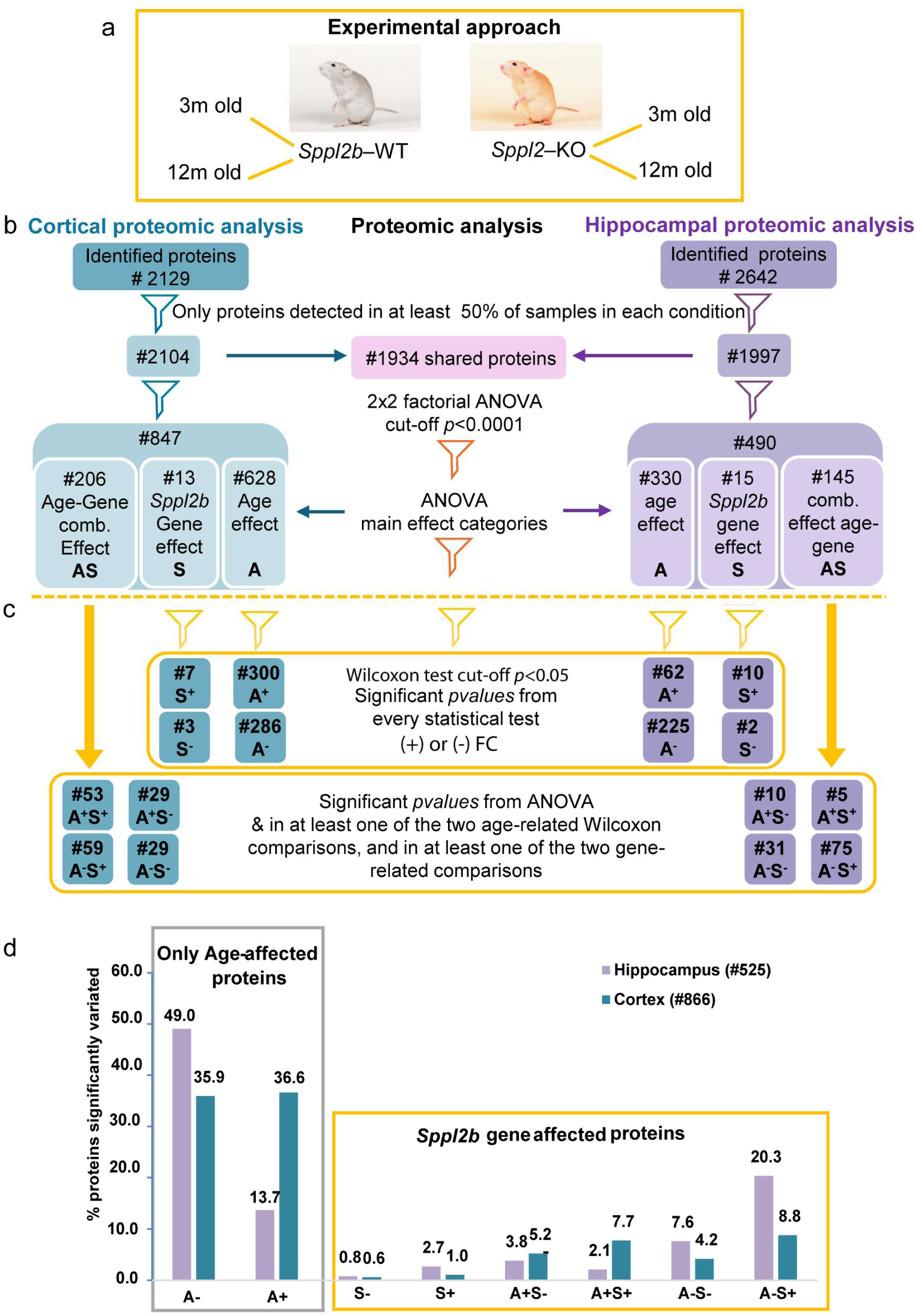
Stratification of proteomic alterations driven by age and Sppl2b deletion in the mouse brain. (a–c) Proteomic data filtering and categorization of protein changes based on statistical parameters obtained by analysis of proteomic quantitative data in cortex (water green) and hippocampus (violet) from *Sppl2b*–KO and WT mice. (b) ‘A’, proteins affected only by age (differences between 3– and 12–month–old mice with ANOVA *p-values*<0.0001); ‘S’, proteins affected only by *Sppl2b* gene deletion (difference between KO and WT with ANOVA *p-values*<0.0001); ‘AS’, proteins affected by both age and the *Sppl2b* gene deletion and with significant changes in relation to both factors (ANOVA *p-values*<0.0001 for at least one of the two factors, and p<0.05 for the other). (c) subcategories of proteins based on fold change (FC) and *p-value p*<0.05 from Wilcoxon tests: A^+^ and A^−^ indicate proteins that increased or decreased with age; S^+^ and S^−^ represent proteins that increased or decreased in KO mice *vs* WT mice; A^+^S^+^, A^+^S^−^, A^−^S^+^, and A^−^S^−^ proteins that changed based on both age and *Sppl2b* gene combined effect. (d) Percentages of proteins identified within each of the 8 subcategories, A^+^, A^−^, S^+^, S^−^, A^+^S^+^, A^+^S^−^, A^−^S^−^, A^−^S^+^, in cortex (water green) and hippocampus (violet).

Whole proteome analysis immediately revealed a dominance of age–driven effects: A^+^ and A^−^ proteins, showing increased or decreased levels with age respectively (Source Data SD1, SD2), represented for more than 60% of all significantly altered proteins in both cortex and hippocampus (Fig. 1d). In contrast, the effects attributable solely to the *Sppl2b* genotype were minor, as the S^+^ and S^−^ proteins, showing increased or decreased levels in KO mice *vs* WT mice respectively (Source Data SD3), were less than 3% of the selected proteins in both the cortex and the hippocampus (Fig. 1D). Proteins that changed based on a combined effect of both age and *Sppl2b* genotype (A^+^S^+^, A^+^S^−^, A^−^ S^+^, A^−^S^−^) showed a broad range of values, but never exceeding 20.3% (Fig. 1D). Notably, the proteomic profile appeared differently regulated in the two brain regions. The percentages of A^+^ and A^−^ proteins were similar in the cortex samples (Fig. 1D, green bars) whereas in the hippocampus the percentage of A^−^ proteins were more than three times that of A^+^ proteins (Fig. 1D, purple bars), indicating a distinct brain region–specific aging signature. Furthermore, the hippocampus and cortex shared 91 A^−^ and 29 A^+^ proteins (Appendix Fig. S1A). Only 1 S^+^ protein, but no S^−^ proteins, was common between cortex and hippocampus (Appendix Fig. S1B). Among the combined–effect subcategories, the cortex and hippocampus shared 1 A^+^S^−^ protein, 2 A^+^S^+^, 4 A^−^S^−^ and 12 A^−^S^+^ proteins (Appendix Fig. S1C). Supplementary Table S1 lists the proteins common between the two brain regions in each subcategory.

Proteomic results for both hippocampus and cortex are presented in the following subsections, organized by the three main effects: i) age, ii) *Sppl2b* gene expression, and iii) their combined effect.

### Proteomic Aging–Dependent Changes in Cortex and Hippocampus of Wild–Type and SPPL2b– Deficient Mice

Age-related trend in protein levels for both A^−^ and A^+^ proteins were similar between WT and *Sppl2b–*KO mice in both tissues, as shown by the correlation of FC values (Appendix Fig. S2 and S3a–d). The application of ANOVA filter (*p<0.0001*) revealed 628 proteins with highly significant aging– dependent changes in the cortex and 330 in the hippocampus samples (Fig. 1b). For downstream analysis, only proteins with significant *p*values in all age–related Wilcoxon tests were considered. This stringent approach yielded a list of age–dependent proteins: 300 A^+^ and 286 A^−^ in the cortex, and 62 A^+^ and 225 A^−^ in the hippocampus (Fig. 1C, Source Data SD1, SD2).

The overall distinction between cortex and hippocampus proteomic profiles was underlined also filtering the proteins with the highest variations with the cut–off Log_2_FC ≥ ±1.5 and – Log_10_*p*values>1.3 obtained by Wilcoxon tests (Table 1). The greatest age–dependent variations, visualized in Volcano plots (Appendix Fig. S4a–d), highlighted key proteins of the aging proteome (Table 1), among which those with high–magnitude changes (Log_2_FC ≥ ±3) are highlighted in blue and red colors for A^−^ and A^+^ respectively. Eight A^−^ proteins and 3 A^+^ proteins were common between cortex and hippocampus, as fatty acid–binding protein 7 (FABP7), GTPase KRas, lamin–B2 (LMNB2), and membrane–associated phosphatidylinositol transfer (PITM1) in the A^−^ subcategory, and heat shock factor binding protein 1 (HSBP1) in A^+^ subcategory. The remaining other age– dependent proteins showed the strongest variations in only one of the two brain regions.

**Table 1.**
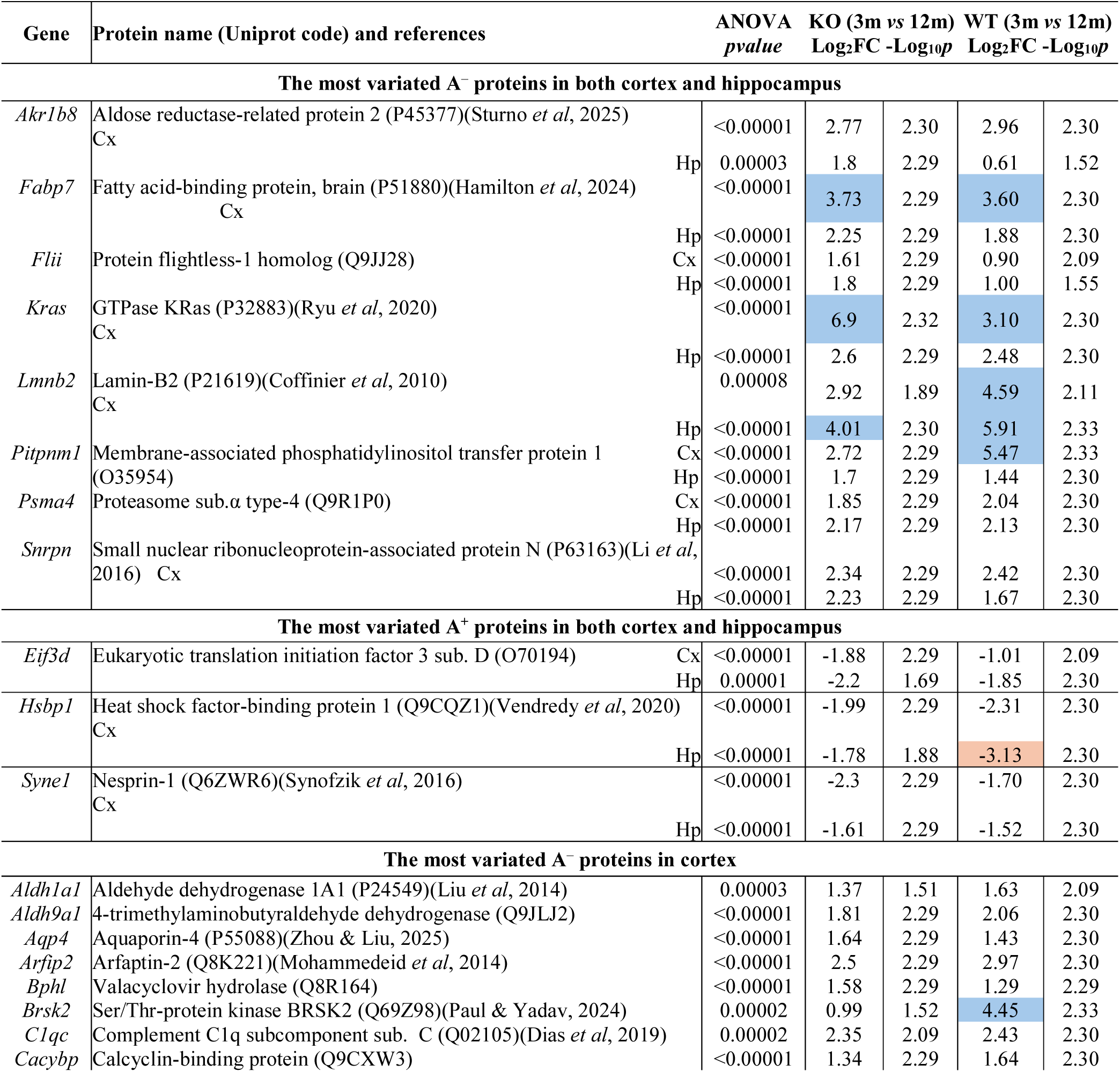
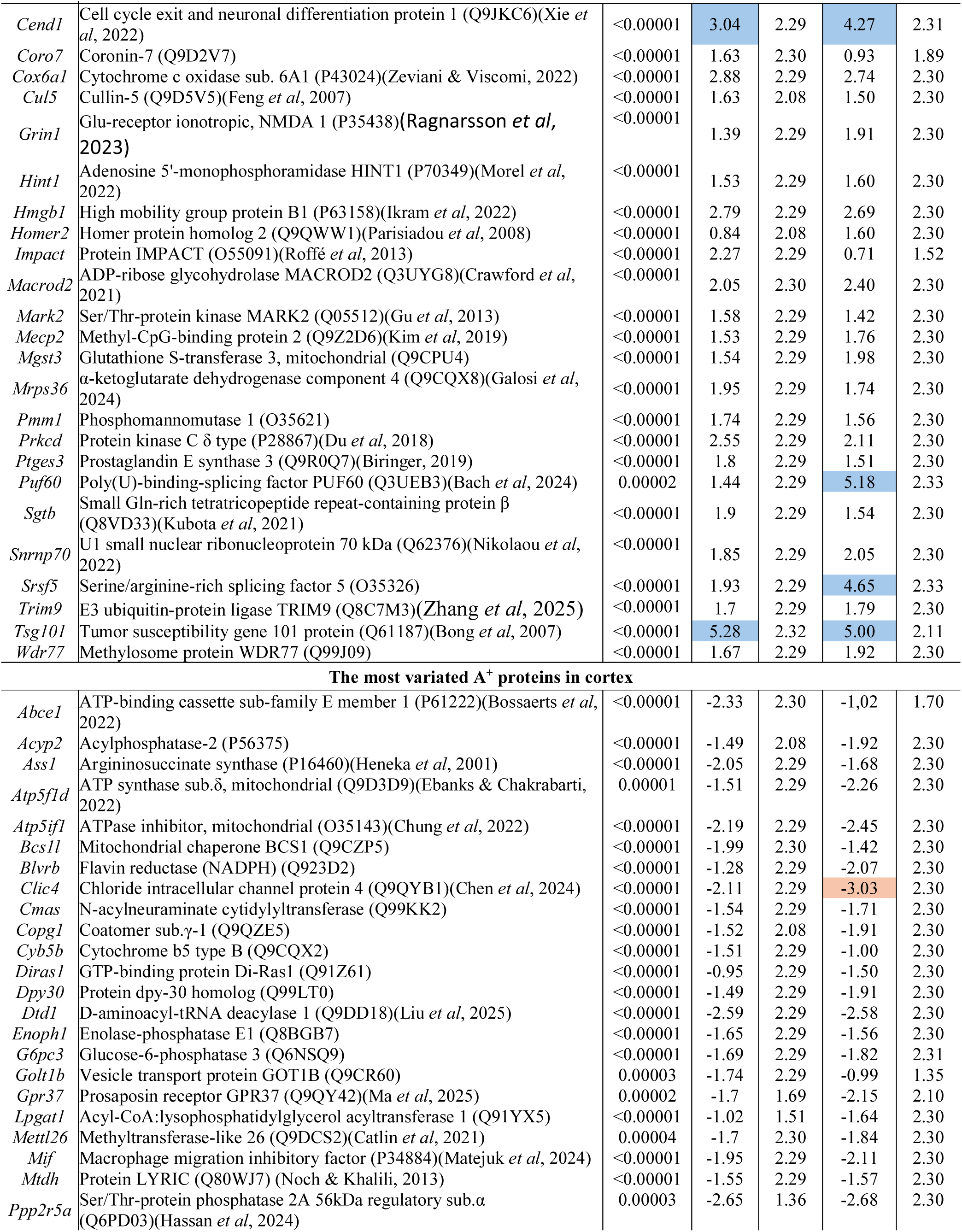
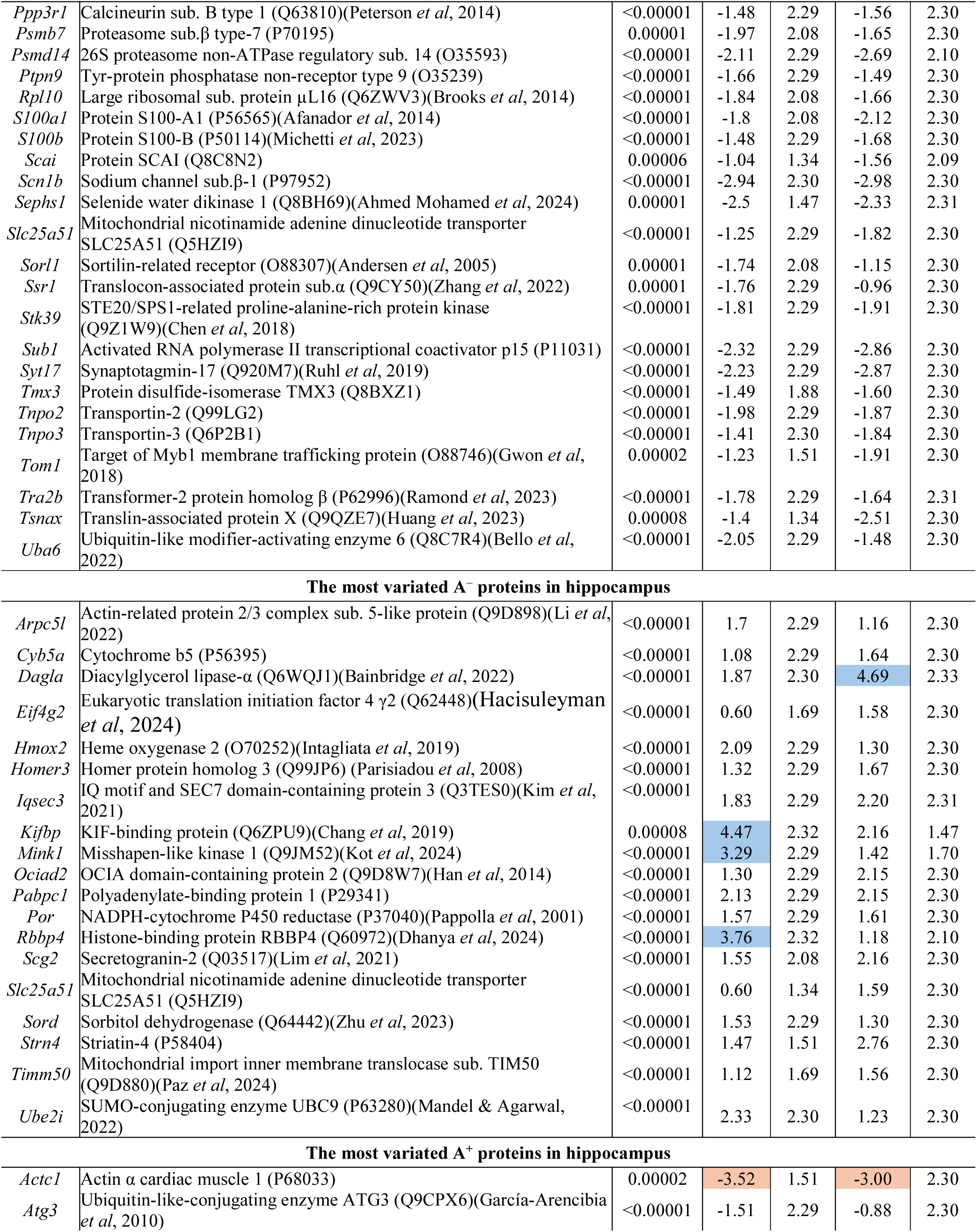
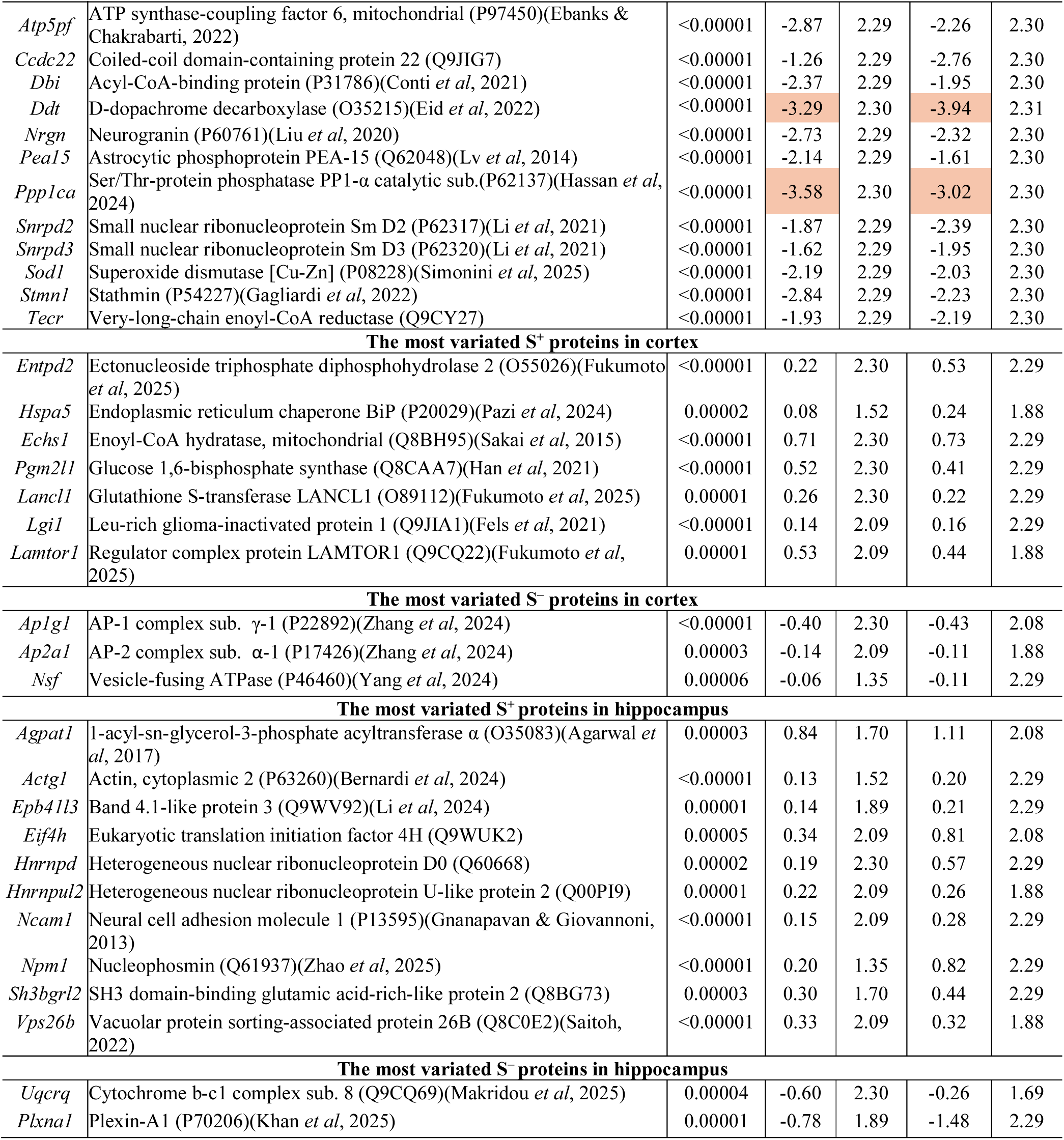
Cortex and hippocampal KO and WT proteins with the greatest variations in relation to a single effect (either aging or *Sppl2b*–genotype). In case of age–dependent variations, listed proteins passed the cut–off Log_2_FC≥±1.5 in at least one of the two model mice, and −Log_10_*p*≥1.3 in each Wilcoxon test. In case of *Sppl*2b–genotype, listed proteins passed the cut–off −Log_10_*p*≥1.3 in each Wilcoxon. *p-*values from nonparametric 2×2 factorial ANOVA, Uniprot codes, and genes are also indicated. Log_2_FC≥±3 are highlighted (blue and red colors for A^−^ and A^+^ subcategories respectively). Cx: Cortex; Hp: hippocampus.

For instance, cell cycle exits and neuronal differentiation 1 (CEND), tumor susceptibility gene 101 (TS101), and serine/arginine–rich splicing factor 5 (SRSF5), were included within the most downregulated proteins specifically in cortex samples. On the other hand, chloride intracellular channel 4 (CLIC4) was the one with the highest FC among the most upregulated proteins (A^+^) (Table 1). The most age–dependent downregulated proteins in hippocampus included diacylglycerol lipase– α (DGLA), kinesin family binding protein (KBP), RB binding protein 4 (RBBP4) and misshapen– like kinase 1 (MINK1). While among the 14 most altered A^+^ hippocampal proteins, actin α cardiac muscle 1 (ACTC), D–dopachrome decarboxylase (DOPD) and Ser/Thr–protein phosphatase PP1–α catalytic sub. (PP1A) showed the highest FC in both KO and WT mice (Table 1). Furthermore, some of the proteins with the most significant age–related variations were found to be in different subcategories in the two brain regions. For instance, high mobility group protein B1 (HMGB1), and homer homolog protein 2 (HOME2), classified as A^−^ proteins in cortex, were found as A^−^S^−^ proteins in hippocampus. Cytochrome b5 type B (CYB5B) belonged to the A^+^ subcategory in the cortex, but it was classified as A^+^S^+^ in the hippocampus. Conversely, several proteins categorized as A^−^ proteins in hippocampus, as heme oxygenase 2 (HMOX2), NADPH–cytochrome P450 reductase (P450R), and secretogranin–2 (SCG2), fell within A^−^S^−^ subcategory in cortex. Finally, ubiquitin– like–conjugating enzyme ATG3 (ATG3) and small nuclear ribonucleoprotein Sm D2 (Sm-D2), classified as A^+^ proteins in hippocampus, were found as A^+^S^+^ proteins in cortex.

### Enrichment analysis of biological processes associated with aging

The enrichment analysis to identify the biological processes (BPs) associated with A^+^ and A^−^ proteins utilized the filtered A proteins (ANOVA *pvalue*<0.0001), from both hippocampus (Source Data SD1) and cortex (Source Data SD2), which were submitted to ClueGO analysis (Appendix Fig. S5–S8, Source Data SD6, SD7). The analysis revealed extensive and distinct functional networks associated with the proteome of the two brain regions (Appendix Fig. S5 and S7, for cortex and hippocampus, respectively), yielding 46 functional GOGroups in cortex and 35 functional GOGroups in hippocampus named with the terms of the most significant BP contained in the GOGroup (Source Data SD6, SD7). The average distribution of A^+^ and A^−^ proteins within functional GOGroups is detailed in Appendix Fig. S6 (cortex) and Fig. S8 (hippocampus), further explicated in Appendix Tables S2 and S3, respectively.

### Proteomic Analysis Reveals a Remodeling of the Cortical Synaptic and Neuronal Proteome During Aging

The majority of GOGroups obtained from cortical proteins by proportions of A^+^ and A^−^ proteins changing similarly from 30% to 70% or *viceversa* (Appendix Table S2, Appendix Fig. S6). Six GOGroups, instead, were associated with a prevalent age-dependent variation (>70% either A^+^ or A^−^). For example, the GOGroup “Regulation of neuron apoptotic process” was predominantly identified by downregulated proteins (79.7% A^−^) (Appendix Table S2, Appendix Fig. S6).

The 10 functional GOGroups identified with the lowest *p*values by the cortical proteins are reported in Fig. 2a, where blue and red bubble size represents the proportion between A^+^ and A^−^ cortical proteins associated with the GOGroups. Fifty–five proteins (24 A^+^ and 31 A^−^), among the most altered in cortex samples, were associated with BPs within the 10 most significant GOGroups (Fig. 2B), indicative of broad functional roles in the biological mechanisms driving cortical aging. The downregulated proteins methyl–CpG–binding protein 2 (MECP2) and glutamate ionotropic receptor NMDA type sub. 1 (NMDZ1) were each involved in seven of the most significant GOGroups. Similarly, the upregulated proteins calcineurin sub. B type 1 (CANB1), S100B, and sortilin–related receptor 1 (SORL) were each associated with five GOGroups (Fig. 2B).

**Figure 2.**
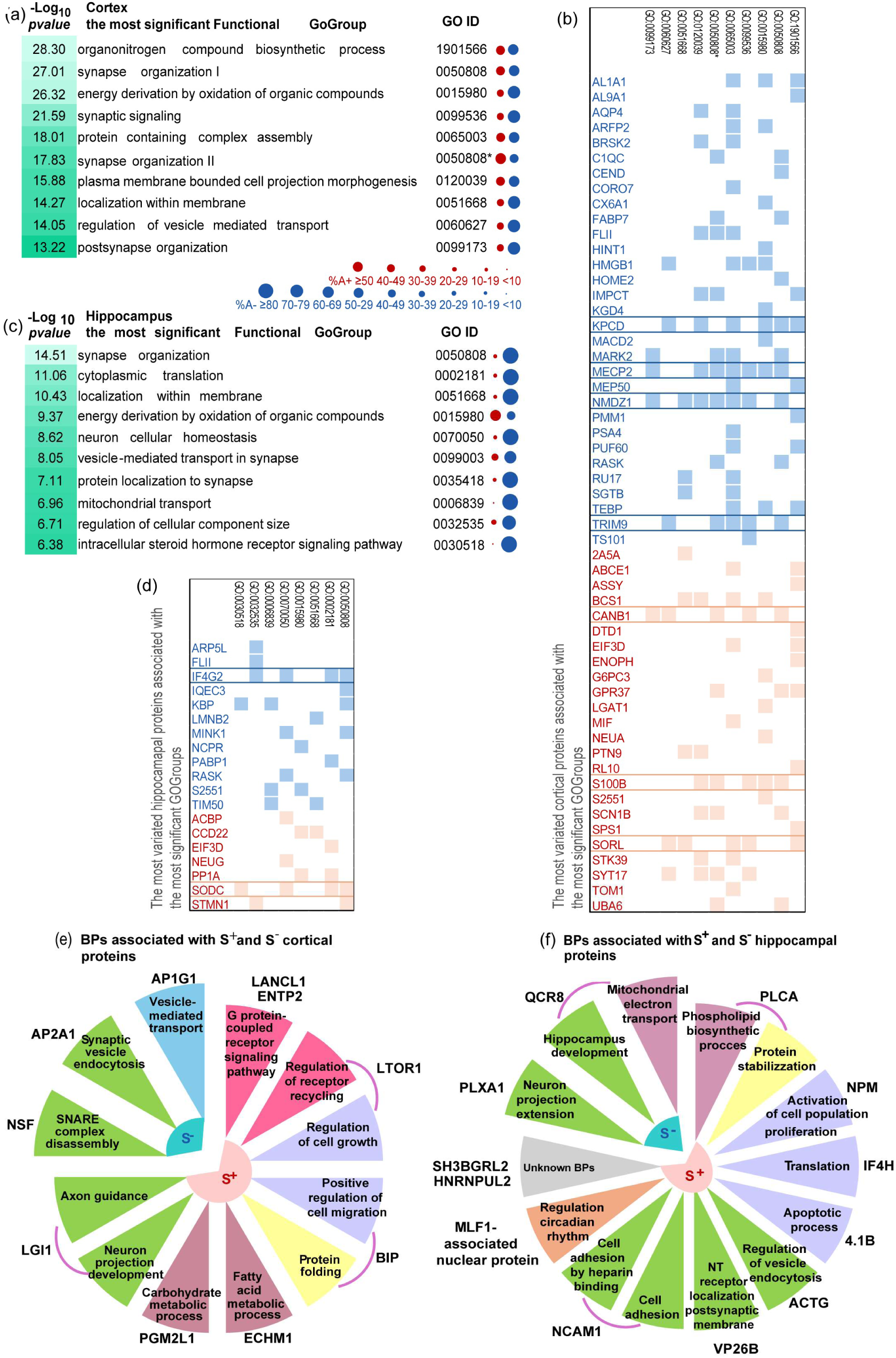
Functional enrichment of age- and Sppl2b-dependent proteomic changes in cortex and hippocampus. (a) The 10 most significant functional GOGroups for cortex proteome listed by decreasing *p-values* order (in −Log_10_ format), relative proportion of A^−^ and A^+^ proteins as represented by the size of red and blue bubbles, respectively. (b) The most variated proteins in cortex samples associated with the 10 most significant GOGroups, red bars for A^+^ and blue bars for A^−^ proteins. The two “Synapse organization” GOGroups are distinguished as I and II in (a) and by an asterisk in (b). The 10 most significant functional GOGroups for hippocampus proteome (c) and the association between the most variated hippocampal proteins and the 10 most significant GOGroups (d). BPs annotated with GO terms for proteins upregulated and downregulated in KO mice (S^+^ and S^−^) in cortex (e) and hippocampus (f).

The most significant GOGroup, “Organonitrogen compound biosynthetic process”, encompassing protein synthesis and translation regulation, was identified by 56% A^−^ and 44% A^+^ associated proteins, which were found to be crucial for neuronal morphogenesis and homeostasis, and shield against neurodevelopmental disorders and neurodegeneration (for references see Table 1). Indeed, this cluster was associated with proteins involved in AD, as SORL, the neuroinflammation–associated prosaposin receptor GPR37 (GPR37), and argininosuccinate synthase (ASSY) that we found to be upregulated, and the downregulated protein kinase C δ type (KPCD). Several downregulated proteins are essential for neuronal development and apoptosis, as the neuronal differentiation regulator IMPACT (IMPCT) (Fig. 2A–B, Table 1). Furthermore, several clusters were centered on core synaptic machinery. The GOGroups “Synaptic signaling” (64% A^−^) and “Postsynapse organization” (70% A^−^) were primarily associated with proteins essential for neurotransmission. These included key neuro–regulators such as NMDZ1 and MECP2, found downregulated, and the senescence– associated CANB1 and the astrocytic marker S100B, found upregulated with aging (Fig. 2B). BPs linked to the synapse organization were grouped in two distinct clusters, indicating functional segregation between BPs related to cognition and glutamate signaling (“Synapse organization I”) and those associated with cell projection and neuron morphogenesis (“Synapse organization II”, Fig. 2A). Structural integrity, cytoskeletal organization and axon extension were substantially affected, with the “Protein–containing complex assembly” GOGroup encompassed among the most significant clusters (Fig. 2A) and associated with the highest number of the most variated proteins (18 A^−^ and 7 A^+^) (Fig. 2B), among which the upregulated SORL, and the downregulated proteins flightless–1 homolog (FLII), KPCD, NMDZ1 and MECP2 (Fig. 2B). BPs included in “Localization within membrane” and “Regulation of vesicle–mediated transport” GOGroups were associated with proteins involved in the regulation of protein targeting and trafficking, exocytosis, synaptic vesicle cycle and regulation of secretory pathways (Source Data SD7).

### Hippocampal Aging is Characterized by a Directional Loss of Synaptic and Translational Capacity

Compared to the cortex, the functional analysis of the hippocampal proteins with age–related variations, revealed a highly directional shift primarily attributable to protein downregulation. Over 70% of the 35 functional GOGroups were characterized by a predominance of A^−^ proteins (Fig. 2C, and, for details, Appendix Fig. S8 and Table S3), with the exception of “Energy derivation by oxidation of organic compounds” and “Axo–dendritic transport” identified by significant percentages of A^+^ proteins (54.4% and 35%, respectively), and “Neurofilament cytoskeleton organization” identified by 100% A^+^ proteins, as superoxide dismutase [Cu–Zn] (SODC), one of the most variated in relation to aging. Contrarily to the cortex, a limited number of proteins, among the most altered (Table 1), were associated with the 10 most significant functional GOGroups and therefore they demonstrated a broad functional influence (Fig. 2D). From this perspective, the downregulated IF4G2 and upregulated SODC, each linked to four of the most significant GOGroups (Fig. 2D), appeared as key proteins in the biological mechanisms underlying hippocampal aging

The most significant functional clusters revealed a pronounced age–related depletion of proteins involved in the core machinery responsible for memory and plasticity, specifically in: (i) “Synapse organization”, (ii) “Cytoplasmic translation”, and (iii) “Membrane and vesicular transport”. The most significant GOGroup was “Synapse organization,” characterized by 88% of A^−^ proteins (Fig. 2C, Appendix Fig. S8 and Table S3). This group included key neuronal processes, such as neuron differentiation, axogenesis, dendrite development, and learning (Source Data SD6). Notably, some of the most downregulated proteins associated with this GOGroup were RASK, MINK1, KBP, and eukaryotic translation initiation factor 4γ2 (IF4G2), while among the most upregulated we found SODC and stathmin 1 (STMN1) (Fig. 2D). “Cytoplasmic translation” GOGroup exhibited significant downregulation (89% associated A^−^ proteins, Fig. 2c, Appendix Fig. S8 and Table S3), indicating reduced protein biosynthesis at both pre– and post–synaptic sites. This GOGroup was associated with the down–regulated IF4G2 and the upregulated SODC, and PP1A (Fig. 2D), among the most variated proteins. BPs included within “Neuronal homeostasis” GOGroup, such as apoptosis, regulation of long–term neuronal synaptic plasticity and axonogenesis may be affected by several proteins with the highest upregulation, as acyl–CoA–binding protein (ACBP), neurogranin (NEUG) and SODC, and downregulation as RASK, MINK1 and IF4G2 (Fig. 2D and Source Data SD6). GOGroups including membrane integrity and signaling were also predominantly identified, such as “Localization within membrane” (83% associated A^−^) and “Vesicle–mediated transport in synapse*”* (70% A^−^) (Fig. 2C, Appendix Fig. S8 and Table S3). Beyond synaptic changes, the hippocampus demonstrated further evidence of structural and metabolic alteration, including mitochondrial impairment. The “Mitochondrial transport” GOGroup was nearly unanimous, with 90% associated A^−^ proteins (Fig. 2C, Appendix Fig. S8 and Table S3) as KBP (Fig. 2D). Similarly, the “Regulation of cellular component size” GOGroup (77% associated A^−^ proteins) indicated cytoskeletal disorganization and impaired actin polymerization, both of which are essential for maintaining synaptic morphology and dendritic spine integrity. The distribution of A^+^ and A^−^ proteins within “Energy derivation by oxidation of organic compounds” GOGroup was more balanced (Fig. 2C, Appendix Fig. S8 and Table S3).

### Deletion of SPPL2b evidenced region–specific proteomic modifications associated with neuronal communication and synaptic pathways

In contrast with the extensive age–related changes, the isolated effect of *Sppl2b* deletion on protein abundance was limited under the first stringent filtering criterion (ANOVA *pvalues*<0.0001). Initial analysis identified 13 proteins (9 S^+^ and 4 S^−^) in the cortex and 15 proteins (13 S^+^ and 2 S^−^) in the hippocampus whose levels were altered exclusively by the *Sppl2b* genotype (Fig. 1B; Source Data SD3). The secondary filtering step was implemented to increase confidence, retaining only proteins that exhibited significant differences between KO and WT mice at both 3 and 12 months of age. This refinement resulted in 7 S^+^ and 3 S^−^ proteins in cortex, and 10 S^+^ and 2 S^−^ proteins in hippocampus (Fig. 1C, Table 1). The final protein set is presented in the heatmap in Appendix Fig. S2. Only one protein, heterogeneous nuclear ribonucleoprotein U–Like 2 (HNRL2), was up–regulated in KO mice in both the hippocampus and cortex, but it passed the secondary filtering step only in hippocampus. The absence of additional shared effects underscores the brain region-specific impact of the *Sppl2b* genotype independent of aging. Functional annotation of these small groups of SPPL2b–regulated proteins (Fig. 2E, F) indicated that SPPL2b primarily modulates targeted processes essential for neuronal communication and maintenance.

#### Cortex

Among the upregulated (S^+^) cortical proteins, Leu–rich glioma–inactivated protein 1 (LGI1) is involved in neuronal structure and signal modulation, including axon guidance and neuron projection development (Fig. 2E), while glutathione S–transferase (LANCL1), and ectonucleoside triphosphate diphosphohydrolase 2 (ENTP2) resulted to be involved in G protein–coupled receptor signaling pathway. The signaling processes and neuronal communications implicate also regulator complex protein LAMTOR1 (LTOR1). This protein is also associated with the regulation of cell growth (Fig. 2E), while the regulation of cell migration is associated with endoplasmic reticulum chaperone BiP (BIP), as well as the protein folding. Protein involved in glucidic and fatty acid metabolism were found upregulated in *Sppl2b*-KO mice, as enoyl-CoA hydratase (ECHM1) and glucose 1,6-bisphosphate synthase (PGM2L1) (Fig. 2E), and HNRL2, a ribosomal regulator of protein synthesis.

The downregulated (S^−^) cortical proteins resulted to be involved in synaptic vesicle dynamics, including vesicle mediated transport and synaptic vesicle endocytosis implicating AP–1 complex sub. γ–1 and AP–2 complex sub. α–1 (AP1G1, AP2A1) respectively (Fig. 2E), and disassembly of the soluble Nsf attachment protein receptor (SNARE) complex involving vesicle–fusing ATPase (NSF) (Fig. 2E).

#### Hippocampus

The upregulated (S^+^) hippocampal proteins, associated with neuronal dynamics, were: neural cell adhesion molecule 1 (NCAM1), involved in cell adhesion mechanisms; actin cytoplasmic 2 (ACTG), involved in vesicle endocytosis; vacuolar protein sorting-associated protein 26B (VP26B), implicated in the localization of neurotransmitter receptors to the post–synaptic membrane (Fig. 2F). Band 4.1-like protein 3 (4.1B), initiation factor 4H (IF4H), and nucleophosmin (NPM) are involved in regulatory mechanisms including apoptosis, protein synthesis, and cell proliferation (Fig. 2F), while 1-acyl-sn-glycerol-3-phosphate acyltransferase α (PLCA) was a S^+^ protein involved not only in lipid biosynthesis but also in the proteostasis. The two downregulated (S^−^) proteins, involved in positive regulation of neuronal processes, were: plexin-A1 (PLXA1), implicated in neuron projection extension; and cytochrome b-c1 complex sub. 8 (QCR8), implicated in the hippocampus development beyond the canonical activity in mitochondrial electron transport chain (Fig. 2F).

### Combined Age and SPPL2b-Genotype Effects Define Distinct Proteomic Signatures in Cortex and Hippocampus

This analysis revealed proteins whose abundance was significantly influenced by a combined effect of age and *Sppl2b* genotype. AS proteins were classified into four subcategories based on the direction of change in both age (A) and genotype (S) comparisons (Fig. 1C, Source Data SD4, SD5): 1) A^+^S^−^ (inverse), abundance increased with age but decreased in KO mice (KO<WT); 2) A^−^S^−^ (concordant downregulation), abundance decreased with age and in KO mice; 3) A^+^S^+^ (concordant upregulation), abundance increased with age and in KO mice; 4) A^−^S^+^ (inverse), abundance decreased with age but increased in KO mice (KO>WT).

As shown in the heatmap reported in Fig. 3A, similar patterns were displayed across both hippocampal and cortical samples, and a huge number of proteins were affected by the combination of age and gene expression effects. Several key proteins were identified in the same AS subcategory in both cortex and hippocampus (Table 1). These included clathrin heavy chain 1 (CLH1) in A^+^S; VAMP2 in A^+^S^+^; AP–2 complex sub. sigma (AP2S1) in A^−^S^−^; kinesin–like protein KIF1A (KIF1A) and apolipoprotein E (APOE) in A^−^S^+^. It is particularly noteworthy that VAMP2 is a known substrate of SPPL2b(Ballin *et al*, 2023), and APOE is major genetic risk factor for Alzheimer’s disease(Islam *et al*, 2025), for this reason these proteins were considered proteins of interest in this investigation. Volcano plots (Fig. 3b–i) visualize the maximum variations induced by the combined effect, applying rigorous cut–off (Log_2_FC≥±1.5 and −Log_10_*pvalue* ≥2.0 from Wilcoxon tests). This analysis clarified which of the two effects, age or *Sppl2b*–genotype, was the predominant driver of the quantitative change for specific proteins.

**Figure 3.**
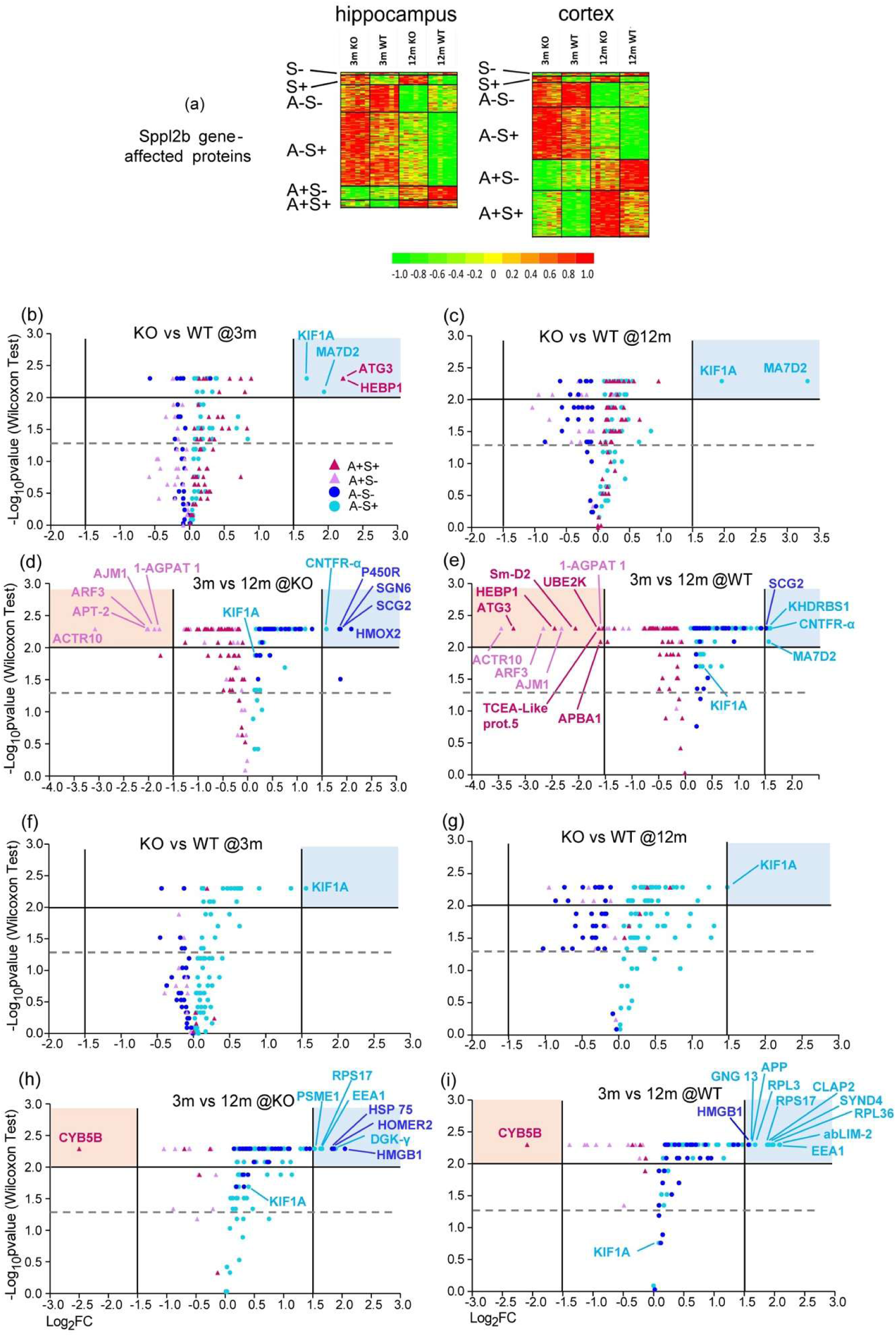
Temporal and genotype-specific modulation of AS proteins in cortex and hippocampus. (a) Heatmap of the proteins showing significant effects (*p*<0.0001) of *Sppl2b* expression in the hippocampus and brain cortex of mice aged 3 and 12 months (3m/12m) and expressing or not expressing the *Sppl2b* gene (WT/KO). Each row of the heatmap represents a specific protein, and each column (six for each group) represents a single mouse. The color of each cell represents the standardized value considering all data for each protein. Shades of green and red represent decreasing and increasing values, respectively, in the range of ± 1 standard deviation. Color tones are accentuated because the heatmap represents only proteins with significant differences. (b-i) Volcano plots visualizing the highest variation of cortex AS proteins in the genotype-related comparison (*Sppl2b*–KO *vs* WT) at 3 months (b) and 12 months (b), and the highest variation in the age-related comparison (3–months *vs* 12–months old mice) for *Sppl2b*–KO (d) and WT (e). The highest variation of hippocampal AS proteins in the genotype–related comparison (*Sppl2b*–KO *vs* WT) at 3 months (f) and 12 months (g) and the highest variation in the age–related comparison (3– months *vs* 12–months old mice) for *Sppl2b*–KO (h) and WT (i). The cut–off is indicated for FC and *p-values* of the Wilcoxon test (1.5, Log_2_FC; 2.0, −Log_10_*pvalue*) evidencing the highest variations. Dotted lines indicate the significance limit (*p*<0.5). Red dots belong to A^+^S^+^ proteins, pink dots to A^+^S^−^, blue to A^−^S^−^, light blue to A^−^S^+^.

#### Cortex

Notably, within A^−^S^+^ subcategory, KIF1A and MAP7 domain–containing protein 2 (MA7D2) resulted to be predominantly driven by *Sppl2b*–genotype effect, since they exhibited markedly the largest FC in both genotype comparisons at 3–months and at 12–months (Fig. 3B,C). In addition, KIF1A significantly varied under the age effect (3–months *vs* 12–months) but not with the highest FC in samples from KO and WT mice (Fig. 3D,E). MA7D2 showed the highest FC also when comparing age groups but only from WT mice (Fig. 3e). Ciliary neurotrophic factor receptor sub. α (CNTFR-α), instead, was highly downregulated with age within A^−^S^+^ subcategory (Fig. 3D,E). The highest variations of proteins within the inverse A^+^S^−^ subcategory appeared to be mainly driven by the age–effect in both KO and WT mice (Fig. 3D,E), among which actin–related protein 10 (ACTR10) and ADP–ribosylation factor 3 (ARF3), in both KO and WT groups, and acyl–protein thioesterase 2 (APT-2) in the KO group (Fig. 3D). For A^+^S^+^ (concordant upregulation) proteins, ATG3 and heme–binding protein 1 (HEBP1) exhibited the prevalent *Sppl2b*–genotype effect in 3– month–old KO mice (Fig. 3d). Within this subcategory, VAMP1 was identified, as well as amyloid-beta A4 precursor protein-binding family A member 1 (APBA1) (Appendix Table S4). APBA1 was one of the AS proteins with the highest FC (Fig. 3E). While, within A^−^S^−^ subcategory (concordant downregulation), the most variated proteins were mainly affected by the aging effect, as SCG2 in both KO and WT mice (Fig. 3D,E) and HMOX2, P450R, and COP9 signalosome complex sub. 6 (SGN6) in KO group (Fig. 3D).

#### Hippocampus

The hippocampus analysis also revealed a higher number of proteins predominantly affected by aging effect, mainly towards a downregulation (Fig. 3F–I). Consistent with the cortex, KIF1A, within A^−^S^+^ subcategory, showed a decreasing of abundance with aging but was also the most upregulated protein in KO mice (Fig. 3F,G). Importantly, the A^−^S^+^ category in KO mice included eleven proteins as small ribosomal sub. protein eS17 (RPS17), early endosome antigen 1 (EEA1), CLIP–associating protein 2 (CLAP2), and APP, showing the highest FC due to a prevalent influence of aging differently to KIF1A (Fig. 3H,I).

The age–related comparisons highlighted significant changes in the A^−^S^−^ (concordant down– regulation) subcategory, where HMGB1 showed the greatest downregulation with age in both KO and WT mice (Fig. 3H,I), suggesting that the age–dependent effect was stronger than the effect of the *Sppl2b* deletion. Finally, the most variated protein within A^+^S^+^ (concordant upregulation) subcategory was CYB5B in both WT and KO mice (Fig. 3H,I).

### Functional Enrichment of Combined Age × SPPL2b Effects in the Cortex

Functional enrichment analysis of AS cortex proteins identified 26 BPs that were grouped into 15 functional GOGroups (Fig. 4A, Source Data SD8). A^−^S^+^ proteins were present in 22 of the 26 biological processes, and they comprised at least 60% of the associated AS proteins in 10 out of the 15 GOGroups, and 100% in 3 GOGroups including “cellular response to amyloid–beta” (light blue, Fig. 4b, Appendix Fig. S10 and Table S5). This finding establishes the A^−^S^+^ subcategory as the predominant functional outcome of the combined effect. A^+^S^+^ proteins were associated with 13 BPs (red color in Fig. 4B, Appendix Fig. S10 and Table EV5) into 8 GOGroups, 3 of which were predominantly associated with A^+^S^+^ proteins only (≥70%) as “membrane assembly” (Appendix Fig. S10 and Table S5). A^+^S^−^ proteins were associated with 16 BPs (pink color, Fig. 4B) within 9 GOGroups and represented 45% of AS proteins in “presynaptic endocytosis” (Appendix Fig. S10 and Table S5). A^−^S^−^ subcategory (blue color in Fig. 4B) consisted in a smaller proportion being associated with 5 BPs into 3 GOGroups among which “canonical glycolysis” where they represented 67.5% of AS proteins (Appendix Fig. S10 and Table S5). Not all the most variated AS proteins contributed to the identification of the functional GOGroups, as evident by comparing Fig. 3B- and Fig. 4C, which reports the specific association to GOGroups of AS protein among those listed in Table EV4. The functional GOGroups defined by the AS proteins were heavily associated with membrane–related processes (Fig. 4A), particularly focused on regulating transport, assembly, and signaling across the cell membrane, as well as the vesicular and axonal dynamics. The most significant functional GOGroup, “presynaptic endocytosis”, was directly associated with VAMP2, A^+^S^+^ protein that contributes to the “synaptic vesicle recycling” BP (Source Data SD8). On the other hand, “Axonal transport” (Fig. 4B,C) was associated with 2 among the most variated AS proteins: ACTR10 (A^+^S^−^) and KIF1A (A^−^S^+^) specifically supporting “anterograde axonal transport” BP. APOE (A^−^S^+^), a major genetic risk factor for AD, was associated with “amyloid–beta formation” GOGroup (specifically with “negative regulation of amyloid–beta formation” BP) and with “cellular component maintenance” GOGroup, including BPs as “maintenance of synapse structure” and “postsynaptic density organization” (Fig. 4A, Source Data SD8). Other prominent GOGroups addressed protein targeting and localization to the membrane, membrane assembly, and the regulation of potassium and sodium ion transport, emphasizing the structural and electrical regulatory functions impacted by the combined AS effect (Fig. 4A). Several functional GOGroups were related to metabolic processes, such as glycolysis, purine ribonucleoside monophosphate processing, and protein catabolism (Fig. 4A), among which, HEBP1 (A^+^S^+^) and HMOX2 (A^−^S^−^), 2 of the most variated cortical AS proteins, were associated with “porphyrin–containing compound metabolic process” (Fig. 4B,C).

**Figure 4.**
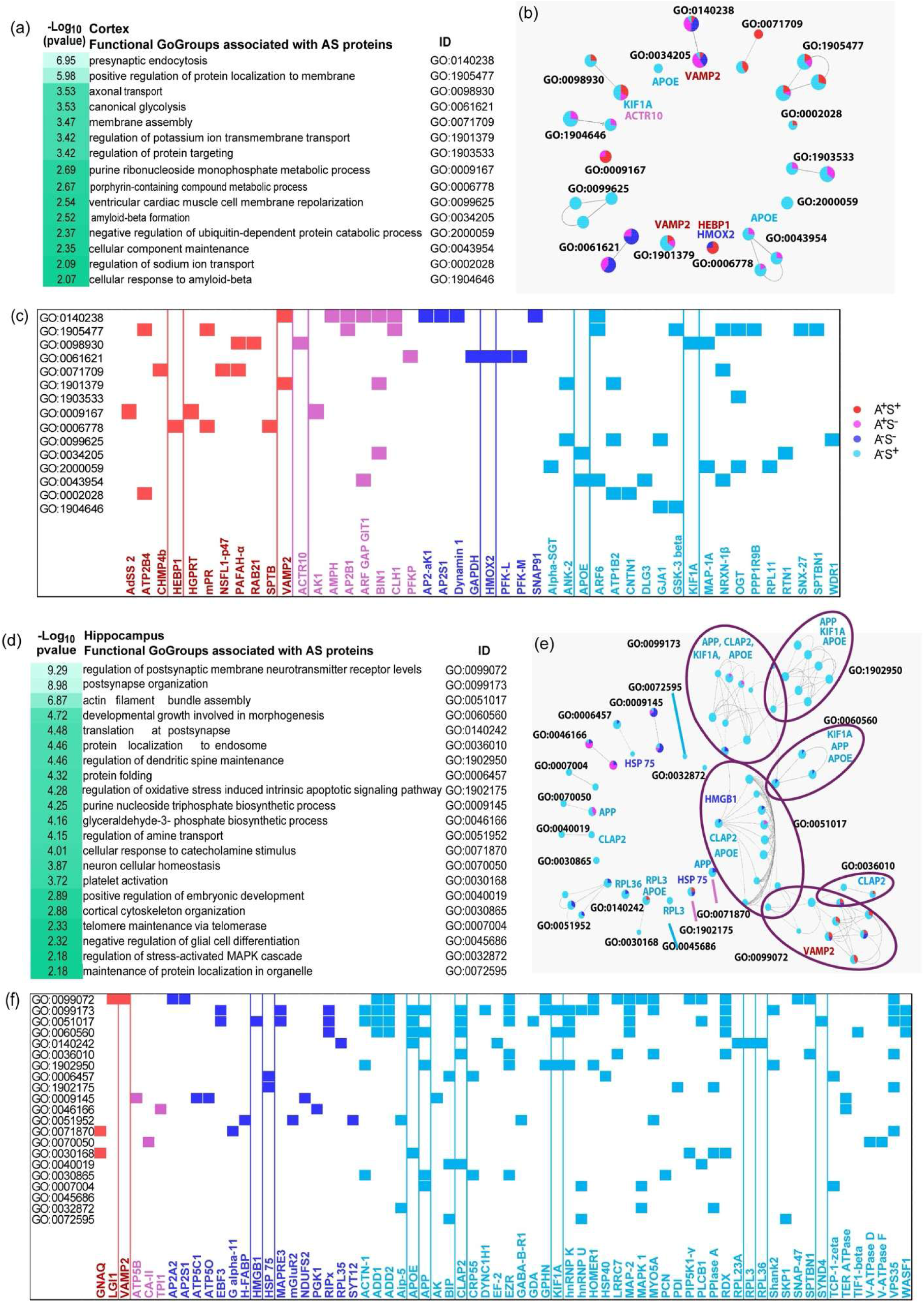
Region-specific functional organization of AS protein networks in the brain. (a) Functional GOGroups provided by ClueGO for AS proteins from the cortical proteome listed in decreasing *p-value* order (in −Log_10_ format) with the GO ID annotations. (b) Functional network identified by cortical AS proteins. Each sphere represents one BP, connected BPs represent a cluster annotated with the ID of the most significant BP, the percentage distribution of AS proteins within each GOGroup is indicated with different colors, red for A^+^S^+^ proteins, pink for A^+^S^−^, blue for A^−^S^−^, sky blue for A^−^S^+^. The most significant AS proteins involved in the GOGroups are indicated. (c) Association between cortical GOGorups and the most variated AS proteins, those evidenced by box are the AS proteins with the highest FC as reported in Fig.3b–e. (d) Functional GOGroups provided by ClueGO for AS proteins from the hippocampal proteome listed in decreasing *p-value* order (in −Log_10_ format) with the GO ID annotations. (e) Functional network identified by hippocampal AS proteins, interconnected GOGroups are evidenced by violet background. (f) Association between hippocampal GOGorups and the most variated AS proteins, those evidenced by box are the AS proteins with the highest FC as reported in Fig.3f–i.

### Functional Enrichment of Combined Age × SPPL2b Effect Targets Hippocampal Synaptic Architecture

Functional analysis of AS hippocampal proteins identified 64 biological processes, which were grouped into 21 functional GOGroups (Fig. 4D, Source Data SD9). The results revealed a distinct and highly directed set of pathways shaped by the combined effect of aging and SPPL2b deficiency. Similarly to the cortical proteome, the A^−^S^+^ subcategory exhibited functional dominance (light blue in Fig. 4E). A^−^S^+^ proteins were associated with 63 up to the 64 identified BPs, within 21 GOGroups, and in 19 of these the A^−^S^+^ proteins represented more than 60% of AS proteins reaching 100% in 8 GOGroups (Appendix Fig. S11, Table EV6). A^−^S^−^proteins (deep blue in Fig. 4E) were associated with 24 BPs and 11 GOGroups, among which “purine nucleoside triphosphate biosynthetic process” (61.9% of A^−^S^−^ proteins) (Appendix Fig. S11, Table S6). The same GOGroup in cortex samples was identified by the prevalent contribution of A^+^S^+^ proteins. In contrast, A^+^S^+^ and A^+^S^−^ proteins contributed minimally, 9 BPs within 4 GOGroups associated with A^+^S^+^ proteins (red in Fig. 4E), 9 BPs within 6 GOGroups with A^+^S^−^ proteins (pink in Fig. 4E). Not all the most variated AS proteins contributed to the identification of the functional GOGroups, as evident by comparing Fig. 3F-I and Fig. 4F. The primary impact of the hippocampal AS effect was on synaptic and structural integrity, specifically synaptic resilience, receptor localization, dendritic spine maintenance, and cytoskeletal dynamics. Indeed, the most significant GOGroups were “regulation of postsynaptic membrane neurotransmitter receptor levels” and “postsynapse organization”. 8 functional GOGroups including neuronal BPs (Fig. 4D) formed a tightly interconnected network centered on high–level neuronal processes (purple ellipses in Fig. 4E). In contrast, many of the remaining clusters were associated with general cellular processes as amine transport, glycolysis, ATP synthesis, and DNA synthesis (Fig. 4D). VAMP2 (A^+^S^+^) was determinant in identifying the most significant GOGroup (GO:0099072), reflecting its role in the localization and transport of neurotransmitter receptors to the postsynaptic membrane (Fig. 4E,F and Source Data SD9). Key structural functions arose from interconnected GOGroups as “regulation of dendritic spine maintenance,” “actin filament bundle assembly,” and “developmental growth involved in morphogenesis” (Fig. 4D). Several AS proteins served as hubs connecting these critical GOGroups, including proteins involved in vesicular and axonal transport, neurodegeneration and resilience, and mitochondrial protection. KIF1A, APP, CLAP2, classified among the most variated A^−^S^+^ proteins in hippocampus proteome, contributed identifying the most significant and interconnected GOGroups a (Fig. 4E,F, Source Data SD9). These proteins participated in multiple regulatory biological processes related to postsynaptic organization, dendritic spine development, and cytoskeleton–dependent intracellular vesicle trafficking. APOE (A^−^ S^+^) was associated with the interconnected GOGroups together with KIF1A, APP, and CLAP2 (Fig. 4E,F, Source Data SD9). CLAP2, APOE and HMGB1 were associated with the regulation of protein– complex assembly (GO:0051017) (Fig. 4D). A^−^S^+^ protein CLAP2 also played a key role in linking protein localization to the endosome, highlighting the relationship between vesicular transport and membrane targeting. APP (A^−^S^+^) specifically contributed to “Neuronal cellular homeostasis” and the “Cellular response to catecholamine stimulation” GOGroups (Fig. 4E,F). Two large ribosomial proteins (RPLs) showed the highest variations within A^−^S^+^ subcategory, the sub. eL36 (RPL36), associated with “Translation at postsynapse” and the sub. uL3 (RPL3), associated with “Negative regulation of glial cell differentiation”, and “Platelet activation” together with APOE (Fig. 4E,F). Heat shock protein 75 kDa (HSP75, A^−^S^−^), a chaperone and heat shock protein, was associated with “Protein folding” and “Regulation of oxidative stress–induced intrinsic apoptotic signaling pathway”.

### SPPL2b Deletion Induces Alterations in Dendritic Spine Density and Neuronal Morphology

Based on functional enrichment data indicating significant alterations in BPs related to synaptic organization and cytoskeletal dynamics in *Sppl2b*–deleted mice, the present study examined whether these changes resulted in quantifiable alterations in neuronal structure.

Golgi staining of 12–month–old *Sppl2b*–KO mice revealed a significant increase in dendritic spine density in both neocortical and hippocampal pyramidal cells, quantified as the number of spines per micrometer (Fig. 5A–C). SPPL2b deficiency was further associated with broader changes in neuronal morphology. Neurons in the KO mouse cortex and hippocampus exhibited significant alterations in soma circularity, area, and perimeter compared to WT controls (Fig. 5D–F). These structural changes in the neuronal soma, consistent with the altered biological processes identified through proteomic analysis being accompanied by significant, quantifiable changes in overall neuronal architecture and cell maintenance. This does not establish that the proteomic changes cause the morphological changes.

**Figure 5.**
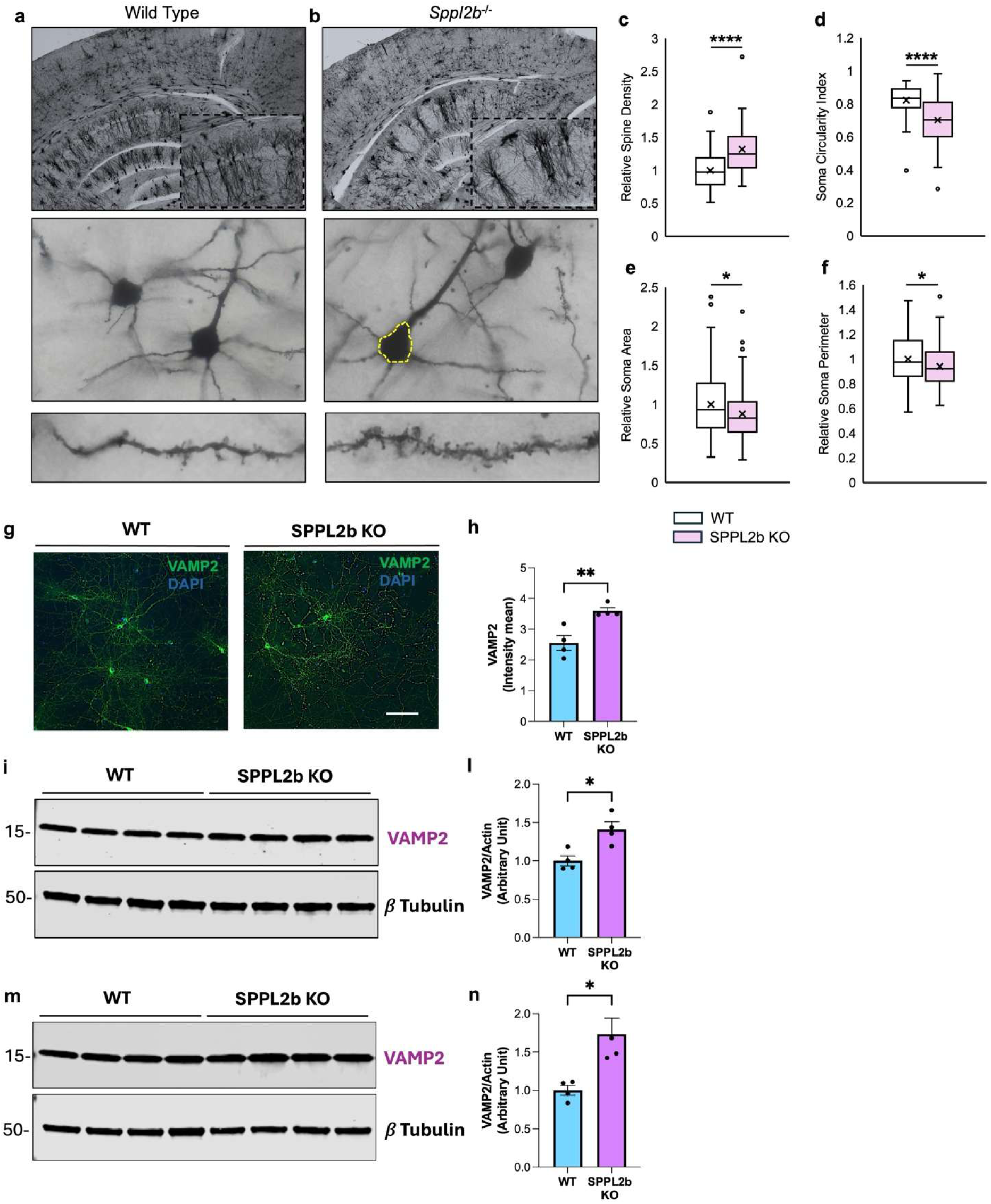
Hippocampal morphology and synaptic alterations in Sppl2b-KO mice. (a, b) Representative Golgi staining images from the hippocampus of 12-month-old Wild Type (a) and Sppl2b-KO (b) mice. Upper panels show low magnification views of the hippocampal region; middle panels show individual neurons at higher magnification; lower panels show dendritic segments with visible spines. Yellow dashed outline in (b) indicates the cell body region. (c) Quantification of relative spine density in Sppl2b-KO compared to WT mice. (d) Soma circularity index measurement comparing WT and Sppl2b-KO neurons. (e) Quantification of relative soma area. (f) Relative soma perimeter measurements. (g) Representative immunofluorescence images showing VAMP2 (green) and DAPI (blue, nuclear stain) in primary neuronal culture from WT and SPPL2b KO mice (Bar Scale= 100µm). (h) Quantification of VAMP2 fluorescence intensity. (i) Representative Western blot showing VAMP2 protein levels in hippocampal lysates from WT and SPPL2b KO mice, with β-Tubulin as loading control. (j) Densitometric quantification of hippocampal VAMP2 protein levels normalized to β-Tubulin. (k) Representative Western blot showing VAMP2 protein levels in cortical lysates from WT and SPPL2b KO mice, with β-Tubulin as loading control. (l) Densitometric quantification of cortical VAMP2 protein levels normalized to β-Tubulin. Data are represented as means ± SEM. Individual data points are shown. *p<0.05, **p<0.01, ****p<0.0001.

Proteomic data strongly implicated VAMP2 as one of the major targets of the combined *Sppl2b*–age effects, notably exhibiting an A^+^S^+^ signature. This finding was validated at both cellular and tissue levels to confirm SPPL2b’s regulatory role in *Vamp2* expression. In primary neurons, immunostaining analysis of VAMP2 in neurons derived from *Sppl2b*–KO mice revealed significantly higher protein intensity at the axonal level compared to WT neurons (Fig. 5G–H). In aged brain tissue, Western blot analysis of 12–month–old mice further validated the proteomic findings, demonstrating significantly higher VAMP2 protein levels in the cortex and hippocampus of KO mice compared to WT controls (Fig. 5I–L).

### Validation of SPPL2b–Dependent KIF1A Regulation

Proteomic data strongly implicated KIF1A, a critical kinesin motor protein involved in axonal transport, as one of the major targets of the combined *Sppl2b*–age effects, notably exhibiting an inverse A^−^S^+^ signature in both cortex and hippocampus. This finding was validated at both the cellular and tissue levels to confirm SPPL2b’s regulatory role in *Kif1a* expression. In primary neurons, immunostaining analysis of KIF1A in neurons derived from *Sppl2b*–KO mice revealed significantly higher protein intensity at the axonal level compared to WT neurons (Fig. 6A,B). These results indicate that the absence of SPPL2b is associated with elevated KIF1A protein accumulation at the neuronal level, particularly within the axonal compartment.

**Figure 6.**
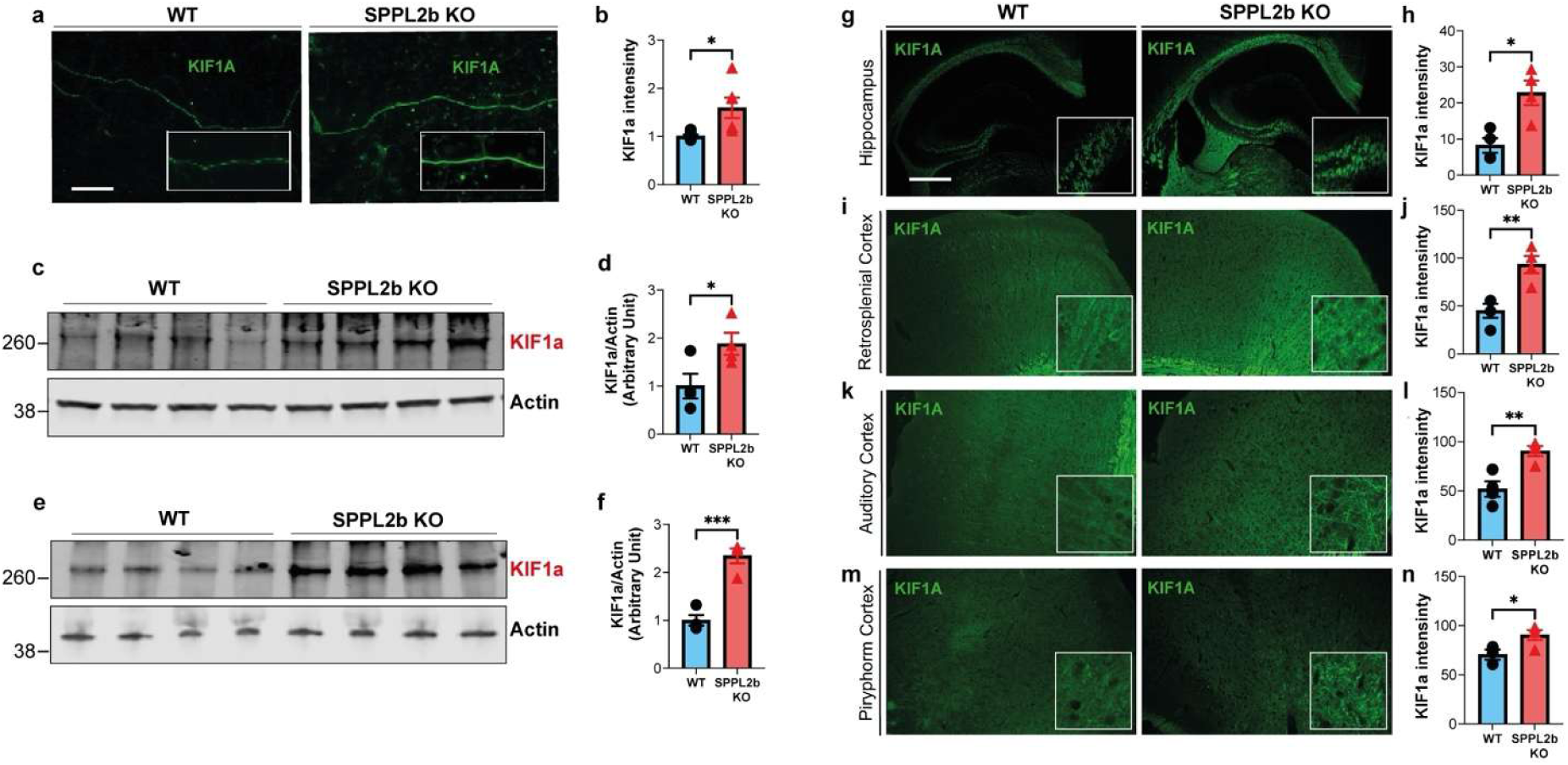
SPPL2b deletion increases KIF1A levels in primary neurons and in multiple cortical and hippocampal regions of adult mice. (a) Representative immunostaining of KIF1A (green) in primary neuronal cultures from WT and *Sppl2b*-KO mice (Bar Scale= 100µm), and (b) corresponding fluorescence-intensity quantification. (c–f) Immunoblot analysis of *Kif1A* expression in hippocampus and cortex from WT and *Sppl2b*-KO mice, with (d, f) band-intensity quantification. Actin was used as a loading control. (g–n) KIF1A immunostaining (green) in brain sections from WT and *Sppl2b*-KO mice, including hippocampus (g, h), retrosplenial cortex (i, j), auditory cortex (k, l), and piriform cortex (m, n), with corresponding quantification. Bar scale =200µm). Data are presented as mean ± SEM. T-test: *P < 0.05; **P < 0.001; ***P < 0.0001.

In aged brain tissue, Western blot analysis of 12–month–old mice further validated the proteomic findings, demonstrating significantly higher KIF1A protein levels in the cortex and hippocampus of KO mice compared to WT controls (Fig. 6C–F). Immunofluorescence analysis also confirmed this protein increase, showing generally higher KIF1A intensity in the hippocampus of KO mice. In the cortex, the increase was particularly pronounced and localized within the retrosplenial, somatosensory, and olfactory areas (Fig. 6).

To determine the underlying mechanism of this protein accumulation, qPCR was performed to assess *Kif1a* gene expression. The results revealed significant upregulation of the *Kif1a* gene expression in the hippocampus of KO mice, with no difference observed in the cortex (Appendix Fig. S11).

### Behavioral phenotyping indicates increased locomotor activity in the absence of significant anxiety–related changes

To evaluate the functional impact of chronic *Sppl2b* deletion, behavioral analyses were performed in 12–month–old female *Sppl2b*–KO mice and WT controls. The Open Field Test (OFT) demonstrated significantly increased locomotor activity in KO mice, as evidenced by greater total distance traveled and higher velocity compared to WT controls (Fig. 7A,B). This elevated activity, which aligns with dysregulation of axonal transport proteins such as KIF1A and MA7D2, indicates that *Sppl2b* deletion may influence general motor function or central arousal. Notably, this increase in movement was not associated with changes in anxiety–related behavior, as time spent and frequency of visits to the central area of the arena did not differ between groups (Fig. 7C,D). To further investigate anxiety– related phenotypes, the Elevated Plus Maze (EPM) and Marble Burying Tests were conducted. In the EPM (Fig. 7E–I), *Sppl2b*–KO mice exhibited behavioral patterns indicative of altered risk assessment rather than increased anxiety. Although KO mice spent less total time in the closed arms (Fig. 7H), the number of visits to this area was unchanged (Fig. 7I). KO mice also spent more time in the central area (Fig. 7G) and visited the open arms more frequently (Fig. 7F), while total time in the open arms was similar to WT controls (Fig. 7E). These findings suggest a subtle shift in exploratory behavior and central area preference, rather than a classical anxiety phenotype. Consistent with these observations, the number of marbles buried did not differ between KO and WT mice, indicating no change in repetitive or anxiety–like digging behavior (Fig. 7K).

**Figure 7.**
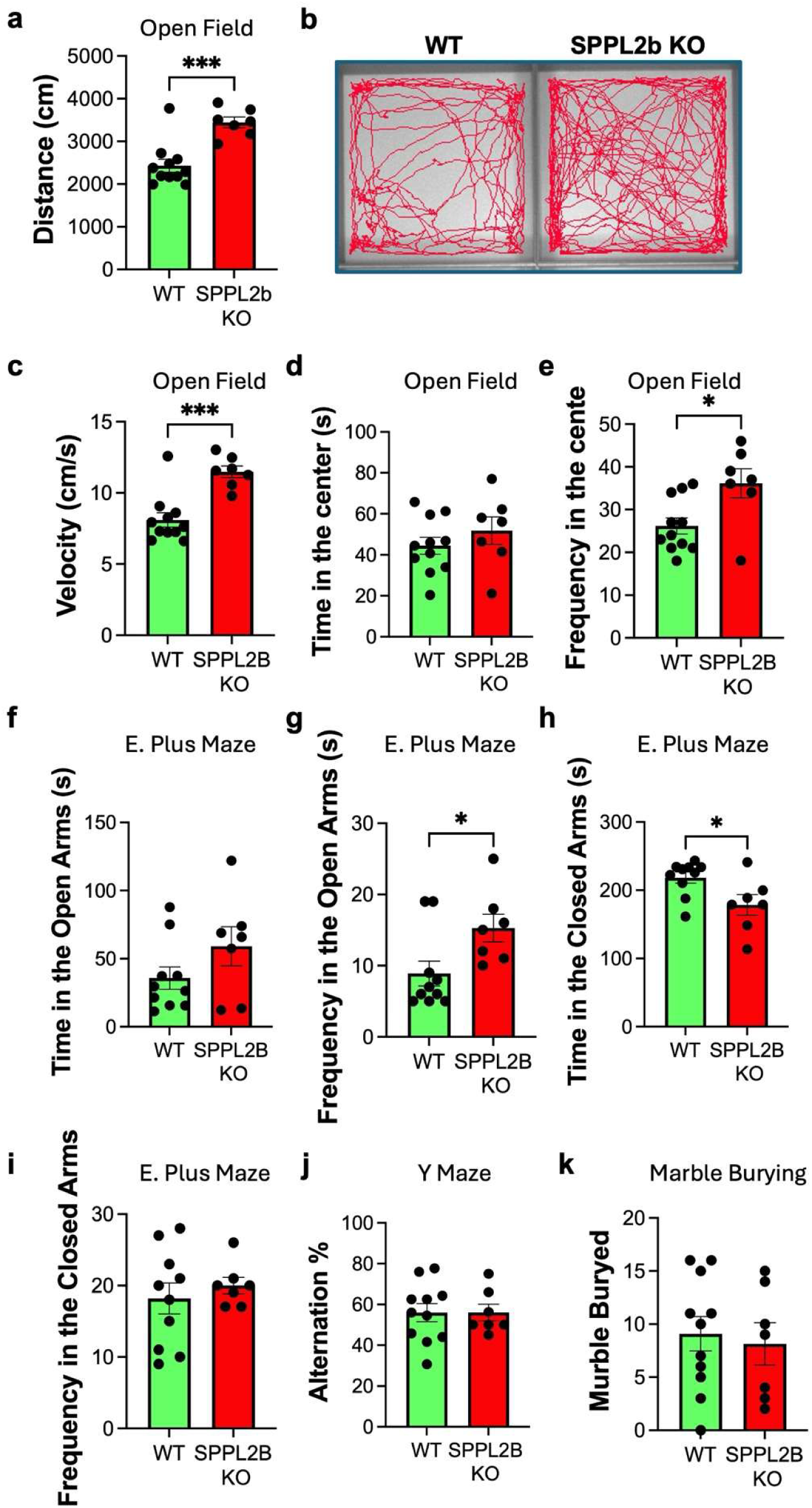
*Sppl2b*–KO mice show increased motility and exploratory behavior. Open field behavioral analysis of total distance traveled (a), velocity (b), time spent in the center of the field (c), and frequency of center visits (d). Elevated plus maze behavioral analysis of time spent in the open arms (e), frequency of entries into the open arms (f), time spent in the center of the elevated plus maze (g), time spent in the closed arms (h), and frequency of entries into the closed arms (i). Y– maze analysis of the percentage of correct alternations while visiting maze arms (j). Total number of marbles buried in the Marble Burying test (k). Data are represented as mean ± SEM. T–test: \**p*<0.05.

The Y–Maze test, which assesses spatial working memory, revealed that the percentage of correct alternations was similar between *Sppl2b*–KO and WT mice (Fig. 7J). These results indicate that, despite observed changes in dendritic spine density and synaptic protein expression, spatial working memory mechanisms remain functionally intact at 12 months of age.

## Discussion

The explorative proteomic profiling of hippocampus and cortex in WT and *Sppl2b–*deficient mice at 3*–* and 12*–*months revealed distinct proteomic alterations associated with aging, *Sppl2b* deletion, and their combined effects. These findings provided important insights on biological processes and brain functional networks affected by SPPL2b deficiency. Behavioral assessments and histological analyses, including Golgi staining and immunofluorescence, provided convergent evidence linking *Sppl2b* deletion to brain biological processes and their networks, particularly synaptic transport mechanisms and neuronal spine and dendritic morphogenesis.

### The aging effect

Proteomic changes induced by aging, in both KO and WT mice, reflected differences in the physiological elderliness of cortex and hippocampus consistent with previous studies (Drulis-Fajdasz *et al*, 2021; Scifo *et al*, 2025). Regional specificity highlighted marked differences in functional networks of the two tissues: cortical up and downregulated proteins in relation with aging contributed similarly to identify functional networks, suggesting a balanced state of decline alongside potential compensatory upregulation. In contrast, hippocampal BPs were predominantly identified by downregulated proteins. The most significant functional clusters associated with cortical proteome revealed alterations of protein synthesis and metabolic processes, extensive remodeling of neuronal structure and synaptic communication. Regarding hippocampal proteome, the functional clusters including synapse organization, neuron differentiation, axogenesis, dendrite development, memory and learning was the most significant. w Notably, about 75% of the most aging–dependent variated proteins identified in this study, were previously associated with neurogenesis, synaptic maturation and neurodevelopment, neuronal homeostasis and metabolism, amyloid–β and Tau processing, and with the control of cognitive and neurological functions. In addition, some of these proteins, exhibiting a wide functional implication in several GOGroups, appeared as critical network nodes in the aging brain, for instance, MECP2 and NMDZ1 (both A^−^) and CANB1, S100B, and SORL (all A^+^) in cortex, while IF4G2 (A^−^) and SODC (A^+^) in hippocampus. The downregulation of MECP2 and NMDZ1 was coherent with their roles in neurogenesis(Kim *et al*, 2019) and in neurodevelopment(Ragnarsson *et al*, 2023). Several cortical proteins involved in neuronal homeostasis, as SPS1(Ahmed Mohamed *et al*, 2024), RL10(Brooks *et al*, 2014) and TRA2B(Ramond *et al*, 2023) were upregulated with aging, probably due to a greater need for homeostatic preservation of cortical cells in older age. It is remarkable that pathological variants of these three proteins were associated with neurodevelopmental deficits in humans (Ahmed Mohamed *et al*, 2024; Brooks *et al*, 2014; Ramond *et al*, 2023). IMPACT protein supports neuronal differentiation under stress by stimulating neurite outgrowth(Roffé *et al*, 2013), and KGD4 affects neuronal metabolism(Galosi *et al*, 2024), both proteins were downregulated in cortex suggesting an aging–dependent physiological decline of their activities. Neuroinflammatory markers such as AL1A1(Liu *et al*, 2014), S100B(Michetti *et al*, 2023), and MIF(Matejuk *et al*, 2024) were differentially regulated with age in cortex, as well as proteins associated with senescence and neurodegenerative pathways as KPCD(Du *et al*, 2018), MARK2(Gu *et al*, 2013), ASSY(Heneka *et al*, 2001), GPR37(Ma *et al*, 2025) and SORL(Andersen *et al*, 2005). Overall, cortex aging appeared associated with both risk factors for neurodegeneration and compensatory neuroprotective mechanisms. For instance, the upregulation of ASSY in older mice appeared to be a risk factor that address towards an improved nitric oxide synthesis and L–arginine metabolism, conditions recognized in the context of AD(Heneka *et al*, 2001). On the contrary, a neuroprotective effect was suggested by several proteins. MARK2, a kinase responsible for Tau phosphorylation in AD(Gu *et al*, 2013), was downregulated in older mice. At the same time, SORL, which is a trans–Golgi sorting receptor for APP, controlling its intracellular trafficking and processing(Andersen *et al*, 2005), was upregulated. The downregulation of FABP7 in both tissues suggested a common mechanism of shutdown of the inflammatory response, outcome in contrast with previous findings demonstrating brain FABP7 upregulation in adult mice(Hamilton *et al*, 2024). Finally, the upregulation of HSBP1 in both tissues is consistent with its protective role against the onset of AD, indeed, it can regulate Aβ oligomerization and Tau fibril formation(Vendredy *et al*, 2020).

Regarding hippocampal proteome, the downregulation of IF4G2 suggested an aging–dependent slowdown in protein biosynthesis necessary for hippocampal homeostasis, synaptic plasticity and long–term memory. This protein can rapidly affect neuronal activity by reprogramming dendritic translation(Hacisuleyman *et al*, 2024). The upregulation of SODC may represent a defense against oxidative stress conditions associated with aging. The downregulation of proteins involved in neurogenesis, as RBBP4(Dhanya *et al*, 2024), and in neurodevelopment, as KBP(Chang *et al*, 2019) and SCG2(Lim *et al*, 2021), suggested their foremost roles in younger age. Analogously, RASK downregulation in older mice was consistent with its important role in hippocampal–dependent learning and synaptogenesis during development(Ryu *et al*, 2020). RASK was downregulated in both tissues, suggesting that its activity is crucial for the development of different brain regions.

### The SPPL2b deficiency effect

Changes specifically induced by *Sppl2b* deletion indicated either probable neuroprotective effects or pathological risk conditions for neuronal development and stability, depending on the dysregulation of specific key proteins. Upregulation observed in cortex for BIP appeared as a neuroprotective effect of the SPPL2b deficiency, in accord with its anti–inflammatory property(Pazi *et al*, 2024), as well as the hippocampal upregulation of NPM, previously proposed as shield against injury–induced stress in brain and as a promoter of neuron survival(Zhao *et al*, 2025). On the contrary, upregulation observed for NCAM1 in hippocampus may negatively affect synaptic adhesion and long–term potentiation(Gnanapavan & Giovannoni, 2013), and that one of E41L3, an apoptosis promoter and AD risk factor(Li *et al*, 2024), supported neurodegenerative vulnerability linked to SPPL2b signaling. Moreover, deletion of the *Sppl2b* gene appeared to affect critical brain processes in both tissues like the endosomal recycling network at synaptic level and consequently synaptic plasticity, and the development of neuron projections and their networks. These were identified by different proteins changing in opposite directions in cortex and hippocampus from KO mice, for instance, LGI1 (S^+^ in cortex) may regulate axon guidance, excitability and synaptic transmission(Fels *et al*, 2021), while PLXA1 (S^−^ in hippocampus) affects neuronal connectivity and anxiety–like behaviors (Khan *et al*, 2025). This is consistent with the altered risk-assessment behavior we observed in the Elevated Plus Maze of 12-month-old *Sppl2b*-KO mice, pointing hippocampal downregulated PLXA1 as a potential molecular candidate correlating with the behavioral shift. The downregulation of NSF, AP2A1, and AP1G1 in cortex could represent a protective mechanism induced by SPPL2b deficiency involving synaptic vesicle recycling and neurotransmitter release. NSF interacts with multiple neurotransmitters and receptors, including SNARE, and its dysregulated/abnormal activity was proved in various neurological and neurodegenerative disorders(Yang *et al*, 2024). AP2A1 and AP1G1 are adaptor proteins (AP) involved in receptor internalization at the postsynaptic terminal, and in the protein trafficking toward the somato–dendritic domain(Zhang *et al*, 2024), respectively. AP may be associated with neurological diseases(Zhang *et al*, 2024), and AP2A1 knockdown reverses senescence–associated phenotypes(Chantachotikul *et al*, 2025). Conversely, the hippocampal proteome in *Sppl2b–*KO mice was characterized by upregulation of proteins involved in the endosome system that supported enhanced postsynaptic recycling, as VP26B(Saitoh, 2022), and movement of synaptic vesicles through the cytoskeletal remodeling, as ACTG(Bernardi *et al*, 2024). Furthermore, SPPL2b deficiency appear to have an effect on metabolism, inducing upregulation of enzymes working in carbohydrate and fatty acid metabolic pathways in cortex, as PGM2L1(Han *et al*, 2021) and ECHM(Sakai *et al*, 2015), as well as in lipid biosynthesis in hippocampus, as 1-AGPAT 1 (Agarwal *et al*, 2017). While a downregulation in hippocampus of QCR8 may associate with mitochondrial dysfunction(Makridou *et al*, 2025). Metabolic alterations disrupt neuronal function and contribute to impair neurodevelopment and to induce neurodegeneration, often resulting from mutations, dysregulation, or deficiency of essential metabolic enzymes(Han *et al*, 2021; Sakai *et al*, 2015; Agarwal *et al*, 2017). SPPL2b could be engaged in networks impacting signaling pathways of the gut–brain–immune axis, which is critical for neuronal and immune homeostasis and connected with neurodegeneration driven by oxidative stress and inflammation(Fukumoto *et al*, 2025). This hypothesis was supported by upregulation in cortex from KO mice of LANCL1 and ENTP2 associated with G protein–coupled receptor signaling, and LTOR1 involved in the mTORC1 pathway. We speculate that this shift toward increased carbohydrate, fatty acid and lipid metabolic activity may in part support the greater metabolic demand associated with the increased locomotor activity we observed behaviorally in *Sppl2b*-KO mice, providing a plausible molecular correlation linking the metabolic and behavioral phenotypes. Because behavior was assessed only at 12 months, in a single cohort of female mice, these differences can be attributed to *Sppl2b* genotype rather than to age, but they do not establish whether this behavioral–metabolic relationship changes across the lifespan or in males.

### The combination of age and Sppl2b deletion effects

Several proteins categorized in the AS group showed significant variations depending on combined effects of age and *Sppl2b* deletion, in both cortex and hippocampus. Interesting results concerned AS proteins implied in neurotoxicity and neuroinflammation, among which HEBP1 (A^+^S^+^) and HMOX2 (A^−^S^−^) in cortex. The concordant effects of SPPL2b deficiency and aging determining the upregulation of HEBP1 could promote neurotoxicity by porphyrin metabolites, an over–expression of this protein indeed was associated with AD(Yagensky *et al*, 2019). On the contrary, the downregulation of HMOX2 could represent a compensatory event by decreasing vulnerability to heme neurotoxicity(Rogers *et al*, 2003). The downregulation of HSP75 and HMGB1 (both A^−^S^−^) in hippocampus indicated probable loss of neuroprotection against oxidative stress(Rego *et al*, 2021), but improved defense against neuroinflammation by depletion of HMGB1, which overexpression was observed in AD(Mo *et al*, 2023). However, the principal finding obtained from the combined effect revealed selective modulation of the stability or expression of key proteins involved in vesicular trafficking, cytoskeletal dynamics and axonal transport, among which KIF1A, APOE, VAMP2 and AP2S1 in both tissues, APP, APBA1, MA7D2 and VAMP1 in cortex, EEA1 and CLAP2 in hippocampus. Though functionally correlated, these AS proteins underwent to different dysregulation. VAMP2 (A^+^S^+^) was upregulated by mutually reinforcing concordant effects of aging and *Sppl2b* deletion, which lead to accumulation of VAMP2 due to lacking proteolysis catalyzed by SPPL2b. VAMP2 contributed to the identification of the most significant functional clusters in both hippocampus and cortex, focused respectively on presynaptic endocytosis and regulation of postsynaptic receptor recycling, as expected for this protein(Ballin *et al*, 2023). VAMP2 is also known as synaptobrevin–2, it is a crucial component of the SNARE complex and is essential for the rapid, calcium–triggered fusion of synaptic vesicles with the presynaptic membrane, thus mediating neurotransmitter release. As an integral vesicle protein, its proper regulation is vital for maintaining synaptic function and plasticity. The effect of VAMP2 may be enforced by the upregulation of VAMP1 in cortex. The concordant downregulation of AP2S1 (A^−^S^−^) was coherent with those of the other two AP, AP2A1 and AP1G1, belonging to S^−^ subcategory in cortex. The slowdown of AP activity may impair the intra–cellular vesicle trafficking, neuronal communications and plasticity, nevertheless, it may be considered also a neuroprotective event induced by *Sppl2b* deletion, with or without the aging combined effect. Indeed, the inhibition or deficiency of AP2S1, avoiding APP processing, contributes to mitigating the cognitive decline in AD patients(Wen *et al*, 2023). Consistently, the aging–dependent downregulation of APP appeared inverted toward an accumulation of APP in KO mice, suggesting a lack of APP processing directly connected to SPPL2b deficiency(Behnke *et al*, 2011), and indirectly through the lessening of AP regulatory proteins as AP2S1. APP was one of the most variated A^−^S+ proteins in hippocampus markedly connected to proteins involved in synaptic resilience and plasticity as APOE, CLAP2, KIF1A, VAMP2 and HMGB1. Among them, APOE is considered a risk factor for AD, being able to interact with Aβ and neurofibrillary tau tangles and to provoke neuroinflammation (Parhizkar & Holtzman, 2022), while KIF1A, due to its microtubule motor activity, is crucial for anterograde axonal transport of synaptic vesicle precursors and for neuronal dense core vesicles transport to the dendritic spines and axons(Okada *et al*, 1995). It is remarkable that KIF1A mutations or reduced expression have been linked to cognitive decline and neuronal loss(Lin *et al*, 2025). Within A^−^S^+^ subcategory KIF1A and MA7D2 are evidently connected in their functions, as MA7D2 stabilizes microtubules acting as cofactor for kinesin transport and may control neurite outgrowth(Kikuchi *et al*, 2022). Furthermore, KIF1A and MA7D2 exhibited the greatest upregulation driven by *Sppl2b* deletion, MA7D2 only in cortex while KIF1A in both tissues showing to be a key brain protein in the axonal vesicle trafficking. Moreover, KIF1A may have a key role in the younger age being a protein decreasing with aging.

The dysregulation of KIF1A was consistent in hippocampus with that of other proteins involved in endosome system, beyond VAMP2, also EEA1 and CLAP2 (both A^−^S^+^), in addition to VP26B and ACTG (both S^+^). EEA1 is a marker of early endosomes(Dong *et al*, 2024), CLAP2 is essential for the endosome maturation during their movement along the microtubules(Dubey *et al*, 2015), activity required for axonal transport, elongation or branching(Dubey *et al*, 2015). Disfunctions of endosomal transport and movement are considered critical factors in the pathogenesis of neurodegenerative diseases, although the related mechanisms remain unclear(Dong *et al*, 2024). The upregulation of EEA1, CLAP2, KIF1A and MA7D2 induced by *Sppl2b* deletion may reflect a protective mechanism that mitigates aging-related neuronal decline. These proteins play essential roles in endosome maturation, axon transportation and dendritic spine development, processes that typically deteriorate with age and contribute to neurodegeneration. Their coordinated increase in SPPL2b–deficient mice suggested a compensatory response that might help preserve molecular pathways critical for maintaining memory-related neuronal architecture. However, we emphasize that our data cannot determine whether *Sppl2b* deletion counteracts age-related changes or whether age modifies the effects of SPPL2b deficiency.

Nevertheless, our findings extended the understanding of SPPL2b’s role in neurodegeneration, particularly its potential involvement in AD, and were consistent with previous studies showing that *Sppl2b* deletion reduces Aβ levels and neuroinflammation, which are hallmarks of AD pathology(Maccioni *et al*, 2024).

Findings on KIF1A were representative of other proteins essential for maintaining axonal transport and synaptic function within cortical and hippocampal circuits, which we highlighted in this study. The interplay between SPPL2b, its substrates as VAMP proteins, and KIF1A represents a candidate pathway that may influence synaptic health in aging and disease contexts; whether the KIF1A is a downstream effector of SPPL2b-related signaling has not been tested directly and remains a hypothesis for future work, especially to understand if an indirect and/or compensatory route is activated due to SPPL2b deficiency. Further morphological and immunological experiments were made to complement and validate proteomic data. These validation experiments (morphology, Western blot, qPCR) were deliberately restricted to 12-month-old animals rather than repeated at both ages, because the proteomic data identified KIF1A and VAMP2 as the most relevant proteins involved in synaptic and trafficking pathways, and 12-month-old was the biologically most robust timepoint for neurodegenerative disease onset, such as AD. This choice means our validation data assessed the aged phenotype specifically and did not establish when across the lifespan these changes first emerged.

Golgi–Cox staining revealed a significant increase in dendritic spine density in *Sppl2b*–deficient mice, particularly in pyramidal neurons, suggesting enhanced synaptic connectivity and plasticity(Moser *et al*, 1994). Consistently, immunofluorescence and western blot analysis showed elevated VAMP2 and KIF1A levels in KO cortex and hippocampus, indicating reinforced synaptic vesicle transport and neurotransmitter release. Notably, the pronounced KIF1A upregulation observed at proteomic level in hippocampus reflected enhanced gene expression triggered by *Sppl2b* deletion, further supporting the hypothesis of a compensatory mechanism counteracting aging-related declines in synaptic maintenance. In contrast, the lack of cortical transcriptional KIF1A changes suggested the presence of a post-transcriptional regulation, but also that *Sppl2b* deletion induced brain region specific adaptation leading to KIF1A upregulation in both tissues, even if gene expression resulted unmodified in the cortex. This region-specific effect is consistent with the distinct cortex and hippocampus proteomic profiles evidenced in the present study.

Behavioral tests demonstrated increased locomotor activity and exploratory behavior in *Sppl2b*– deficient mice. While the upregulation of proteins engaged in the synapse organization, such as KIF1A and VAMP2, together with increased spine density, may underline these enhanced behavioral outcomes by promoting synaptic efficiency and neurotransmission, the behavioral-proteomic relationship may be correlated with several features. For instance, the concordant upregulation of metabolic enzymes described above (PGM2L1, ECHM, 1-AGPAT1) may provide the additional energetic capacity needed to sustain the elevated locomotor output. Similarly, several of the S^+^/S^−^ proteins discussed above with roles in receptor signaling and neuronal connectivity, including LGI1, PLXA1 and NCAM1, are independently implicated in anxiety-related circuitry, offering candidate molecular correlates for the altered exploratory and risk-assessment behavior we observed in the Elevated Plus Maze (Kondo *et al*, 2012)(Lin *et al*, 2025)(Moser *et al*, 1994)(Fels *et al*, 2021; Drabek *et al*, 2006)

### Limitations

Several limitations constrain the interpretation of these findings. First, *Sppl2b* deletion in this model is constitutive, so the proteomic and morphological differences observed at 3 and 12 months cannot be fully dissociated from developmental effects of lifelong SPPL2b deletion; an inducible or conditional knockout would be required to isolate effects specific to adult or aged tissue. Second, proteomic profiling was performed on whole cortex and hippocampus, which precludes attribution of the observed changes to specific cell types (neurons, astrocytes, microglia); given that SPPL2b is expressed in both neurons and glia, cell-type-resolved or single-cell approaches will be needed to clarify the cellular origin of these signatures and to determine whether tissue-level averaging masks more pronounced cell-type-specific effects. Third, while we validated selected proteins (KIF1A, VAMP2) at the protein and transcript level and linked their upregulation to changes in dendritic spine density and locomotor behavior, we did not directly test axonal transport, vesicle trafficking kinetics, or synaptic transmission; the proposed SPPL2b–KIF1A relationship to axonal transport and synaptic function is therefore inferred from correlative and structural data rather than established functionally. Fourth, several morphological, biochemical, and behavioral experiments were conducted with modest sample sizes, and behavioral phenotyping was restricted to 12-month-old female mice, which limits generalizability across sex and age.

## Conclusions

This exploratory proteomic study demonstrated that SPPL2b deficiency leads distinct remodeling of the synaptic and axonal transport proteome in cortex and hippocampus. *Sppl2b* deletion was associated with coordinated upregulation of proteins involved in vesicular trafficking and microtubule-dependent transport, including the kinesin motor protein KIF1A, together with increased dendritic spine density and altered locomotor and exploratory behavior. These findings point to a previously unrecognized association between SPPL2b and synaptic proteostasis and neuronal architecture, processes that are disrupted in neurodegenerative disorders. The candidate SPPL2b– KIF1A relationship offers a testable hypothesis linking intramembrane proteolysis to axonal transport and synaptic function. However, causality and cell-type specificity remain to be established. Overall, our results propose new insights into SPPL2b role in modulating axonal transport and synaptic plasticity, key processes in the context of neurodegeneration.

## Methods

### Mice

The study was conducted using mice bred at the Comparative Medicine Biomedicum (KM–B) facility, located on the Solna campus of Karolinska Institutet. Embryos of *Sppl2b*–knock–out (KO) mice (B6; CB–3110056O03RikGt(pU–21T)160Imeg) were sourced from the Center for Animal Resources and Development at Kumamoto University and implanted at the Karolinska Center for Transgene Technologies (KCTT) Comparative Medicine facility. Wild–type (WT) mice with a C57BL/6J background served as controls. The animals were housed in groups of three to five under a 12–hour light/dark cycle (lights on at 7:00 a.m.). All animal experiments were conducted in accordance with the Swedish Board of Agriculture regulations (SVJFS 2019:009, L150) and complied with the European Directive 2010/63/EU for animal experiments. The study was approved by the Stockholm Animal Ethics Committee (15758–2019 and 12570–2021). A total of 23 female mice (with 5 to 6 per group), aged 3 and 12 months, were included in the proteomic study. The animals were anesthetized using 2% isoflurane and perfused intracardially with PBS. The experimental groups comprised five WT and six *Sppl2b*–KO mice at 3 months of age, and six WT and six *Sppl2b*–KO mice at 12 months of age. Following perfusion, the brains were rapidly extracted and divided into two hemispheres. One hemisphere was fixed in 4% formalin for morphological and immunohistochemical analyses, while the other hemisphere was dissected to isolate the hippocampus and cortex, which were promptly frozen at −80°C for subsequent proteomic analyses.

### Samples’ treatment for shot–gun proteomic analysis

Frozen brain tissue samples from all the animals were homogenized in RIPA buffer (ThermoFisher Scientific, Waltham, MA, USA) containing a mixture of protease inhibitors (Mammalian Protease Arrest, G biosciences) in a ratio of 1:100 v/v. The volume of extraction buffer was 300 μL for each hippocampus, and 800 μL for each cortex. Homogenization of samples was achieved in 5 cycles by using a pestle for 20 seconds followed by 20 seconds of rest on ice. Samples were then subjected to 3 cycles of 1 minute sonication in an ice bath, and then centrifuged at 20800 rcf for 20 minutes at 4°C. The supernatant of each sample was recovered, and total protein concentration was measured by bicinchoninic acid assay (BCA, Sigma Aldrich, St. Louis, MO, USA) following the manufacturer’s instructions. Samples were diluted to the same final protein concentration (0.5 μg/μL) in 50 mM TrisHCl pH 8.5, and 50 μL of each sample was submitted to the Filter Aided Sample Preparation (FASP) protocol(Wiśniewski, 2019) occurring into 30 kDa Microcon filters (Merck, Darmstadt, Germany). Highly specific trypsin MS Grade (Pierce™, Thermo Fisher), 0.1 μg/μL in 40mM ammonium bicarbonate pH 7.8/acetonitrile 10:1 v/v, was added at ratio 1:50 w/w enzyme/substrate, and incubation occurred overnight at 37°C. Finally, tryptic digested samples were washed three times by adding 50 uL of 50mM ammonium bicarbonate pH 7.8 followed by centrifugation at 14000 rcf for 15 min. All the samples were then dried and de–salted with OMIX C18 pipette tips (Agilent, Santa Clara, CA United States) following the manufacturer’s instructions. The tryptic peptides were solubilized in 250µL 0.1% formic acid (FA) and immediately stored at −80°C until HPLC–(HR)–MS/MS.

### High**–**resolution mass spectrometry analysis

All chemicals and reagents were of analytical grade and were purchased from Sigma–Aldrich/Merck, Darmstadt, Germany. The HPLC–(HR)–MS/MS apparatus consisted in a HPLC Ultimate 3000 (Dionex, Sunnyvale, CA, USA) equipped with a FLM–3000–Flow manager module and coupled by a nano–electrospray source (nano–ESI) to an LTQ–Orbitrap Elite MS (ThermoFisher Scientific San Jose, CA, USA). The injection volume was 10 µL of each sample randomly ordered, and for chromatographic separation a reversed–phase Easy Spray C18 nano–column was used (250 mm length, 75 µm inner diameter, and 2 µm particle size, Thermo Fisher Scientific). Mobile phases were: A, aqueous 0.1% FA; B, 20% aqueous 0.1% FA/80% acetonitrile (v/v), mixed at 0.3 µL/min flow rate with the following gradient: 4%B for the first 3 min., 50%B at 80^th^ min, 80% B at 90^th^ min., 90% B at 92^th^ min. Full (HR)–MS measures were performed in positive ion mode from 350 to 1600 m/z, resolution 120,000 at 400 m/z, capillary temperature 275°C, source voltage 1.6 kV and S–Lens RF level 68%. The 10 most intense ions were fragmented in data–dependent acquisition mode by collision induced dissociation (35% normalized collision energy for 10 ms, 2 m/z isolation width, activation q of 0.25). (HR)–MS and MS/MS data were generated by Xcalibur 3.0.63 (Thermo–Fisher Scientific). MS/MS fragmentation spectra were analysed by Proteome Discoverer (PD) software (version 2.2, Thermo Fisher Scientific) with the SEQUEST HT cluster search engine (University of Washington, licensed to Thermo Electron Corporation, San Jose, CA, USA) against the UniProtKB mouse database (reviewed, 37,517 entries, release 2023_02). PD settings were: peptide mass tolerance 10 ppm; fragment ion mass tolerance 0.6 Da; 2 missed tryptic cleavages; FDR 0.01 (strict) and 0.05 (relaxed); Cys–carbamidomethylation as fixed modification; Met–/Trp–oxidation, Ser–/Thr–, Tyr–phosphorylation, N–terminal pyroglutamic acid and N–terminal or Lys acetylation as dynamic modifications. Peptides were filtered for high confidence and a minimum length of 6 amino acids. Only proteins identified in at least the 50% of each group (WT, *Sppl2b*–KO, 3 and 12 months– old) have been subjected to PD Label–Free Quantification (LFQ) and normalized against the total peptide amount. Keratins and hemoglobin were considered contaminants and thus excluded. The MS data have been deposited to the ProteomeXchange Consortium via the PRIDE(Perez-Riverol, 2022) partner repository with the dataset identifier PXD060275.

### Statistics and data Filtering

The analysis was performed on cortex and hippocampus brain regions from KO and WT mice aged 3 and 12 months. The experimental design included 4 groups, each composed of 6 mice: 6 WT aged 3–month–old, 6 WT aged 12–months–old, 6 KO aged 3–month–old and 6 KO aged 12–months–old. One 3– month–old WT mouse was lost, so its missing data were replaced with the median of the 5 remaining mice of the same group. To ensure high confidence for the comparison across aging– related (3– and 12–month–old mice) and genotype–related (WT and *Sppl2b*–KO) conditions, only proteins detected in at least 50% of the samples per condition were considered. To assess the effects of age and *Sppl2b* gene deletion, 2×2 factorial ANOVA and Wilcoxon tests were applied using the nonparametric aligned rank transform method(Kay *et al*, 2021).

Significant statistical data were filtered by ANOVA *pvalues*< 0.0001. Log_2_ fold change (FC) was calculated as the difference between the group medians. For the aging effect the Log_2_FC was calculated as the difference between 3– and 12–month–old mice, for both KO and WT. Consequently, positive Log_2_FC was obtained for proteins with higher levels at the age of 3 months, and negative Log_2_FC for proteins with higher levels at the age of 12 months. A^+^ symbol was adopted for proteins increasing with age, and A^−^ symbol for those decreasing with age. For the effect of *Sppl2b* gene expression, the Log_2_FC was calculated as the difference between KO and WT mice, both at 3 and 12 months of age. Thus, proteins more abundant in KO mice resulted in positive Log_2_FC values and denoted by S^+^ symbol, proteins less abundant in KO resulted in negative Log_2_FC values and denoted by S^−^ symbol. The symbol “AS” was applied to those proteins affected by both age and the gene and with significant changes in relation to both factors. ANOVA was followed by single Wilcoxon tests to compare the age– and *Sppl2b*–related effects in two-group comparisons: 3–months *vs* 12–months in WT mice and in KO mice; KO *vs* WT in 3–month mice and in 12–months mice. All tests were repeated for hippocampus and cortex proteins. We considered acceptable statistical data when outputs of ANOVA and Wilcoxon tests agreed, and the proteins showed Wilcoxon *pvalues*<0.05 in both two age–based comparisons (in KO or WT group), or in both two KO vs WT comparisons (either at 3 or 12 months of age) (Source Data SD1–SD3). In the case of AS proteins, data were considered acceptable when significant results (*p*<0.05) were obtained from direct Wilcoxon tests in at least one of the two age-related comparisons (3m vs 12 m in KO and/or in WT), and in at least one of the two gene-related comparisons (KO vs WT 3m and/or KO vs WT at 12m) (Source Data SD4–SD5).

### Functional analysis

Functional analysis was performed by ClueGO plugin (v. 2.5.10) via Cytoscape software (v. 3.10.2)(Mlecnik *et al*, 2018). For both cortex and hippocampus samples, the list of significant proteins varying with respect to age and with respect to both factors, age and *Sppl2b* gene, were imputed on ClueGO and tested for Mouse GO Biological Process (last updated 8th February 2024) using the enrichment right–sided hypergeometric test and the Benjamini–Hochberg correction statistical options. Only biological processes with *p*<0.01 were accepted, with the kappa score set at 0.4. Minimum and maximum tree interval values were 3–8, and a minimum of 3 genes and 4% of genes were selected for GO terms, except for functional analysis of proteins varied in cortex samples based on age, where a minimum of 6 genes and 4% of genes for GO terms was set. Go Term Fusion was selected, while the evidence code decision tree was set to “all”, excluding only Electronic GO annotations (IEA). BPs involving proteins up–regulated and down–regulated in KO mice (S^+^ and S^−^), in cortex and hippocampus, were annotated by the QuickGO tool (https://www.ebi.ac.uk/QuickGO).

### Behavioral analysis

The mouse behavioral tests were conducted using 12–13–month–old female WT and *Sppl2b*–KO mice, in total were used 12 to 7 mice respectively per group. The mice underwent a sequence of behavioral tests in the following order: Open Field (OF) > Elevated Plus Maze (EPM) > Y–Maze > Marble Burying (MB). Experimental procedures were performed under white light (100 lux) from 10:00 to 17:00. Mice were transferred to the experimental room one hour before the start of behavioral testing for acclimatization. The apparatuses were cleaned with 70% ethanol between each mouse. The EthoVision XT 15 video tracking system (Noldus Information Technology) was used for monitoring, recording, and analyzing the mouse behavioral tests. *Open field (OF).* For the OF test, it was used an open square arena with 45 × 45 x 45 cm size. The mice were initially positioned in the center of the arena and allowed to explore the area for 5 minutes. After the 5 minutes the mice were removed and repositioned inside the home cage. Total distance, velocity, time in the center and time spend in the border were obtained. *Elevated plus maze (EPM).* EPM consisted of two closed arms and two open arms located at 40 cm height. Mice were placed in the center of the plus maze and allowed to explore for 5 min. Several parameters were acquired including the number of entries to closed or open arms, time spent in closed or open arms, and frequency of head dips. *Marble burying (MB).* MB tests were performed in a cage with dimensions of 40 x 35 x 22 cm and filled with 5 cm deep fresh wood chips for bedding. 20 glass marbles (5 × 4) were placed evenly, and mice were allowed to explore the marbles for 30 minutes. The number of buried marbles was recorded. *Y*–*maze.* Y–maze consisted of three identical arms in a “Y” shape. Mice were allowed to explore the other two arms for 10 min. Percent alternation is defined as the number of spontaneous alternations/the total possible triads. The percentage of alternation (% alternation) was calculated as {spontaneous alternation/(total number of arm entries - 2)} × 100.

### Golgi staining and dendritic spine analysis

After behavior, we randomly selected 5 female Sppl2b-WT and Sppl2b-deficient mice and processed them for Golgi analysis. We stained 150–μm–thick coronal sections from the left brain hemispheres with the Golgi-Cox solution using the FD Rapid GolgiStain Kit (NeuroTechnologies, USA), following the manufacturer’s instructions. We acquired images from a single focal plane and a merged z–stack with a light microscope (Nikon Eclipse E800) with 100× objective, oil immersion (NA: 1.30). We performed the analysis on neocortical and apical collateral dendrites (stratum radiatum) from CA1 pyramidal neurons. For the analyses, we selected neurons based on the following criteria: (i) neurons relatively complete (two orders or greater dendrites were entirely visible and in focus); (ii) neurons fully impregnated with staining; and (iii) minimal or no overlap with other labeled neurons. We selected 5 to 10 different neurons per brain region per mouse and measured one to two dendrites per neuron for a total dendritic length of at least 300 μm per animal. All analyses were conducted blinded to genotype. Imaging analyses were performed with ImageJ software (National Institutes of Health, USA).

### Western Blot

Mouse brain cortex and hippocampal tissues were lysed using RIPA buffer (ThermoFisher) supplemented with phosphatase inhibitors (phosphatase inhibitor cocktails, Sigma) and protease inhibitors (mammalian protease arrest, G Biosciences). The tissues were homogenized, sonicated, and centrifuged at 14,000 g for 20 minutes at 4°C. The resulting supernatants were collected, and protein concentrations were determined. Samples were stored at −80°C until further use. Protein concentrations were quantified using the Pierce™ BCA Protein Assay kit. For Western blot analysis, 20 µg of protein per sample was loaded per well and normalized to β–actin and β–tubulin. The samples were loaded in Mini–PROTEAN® TGX™ Precast Gels (4–20%, Bio–Rad) and transferred onto nitrocellulose membranes (Bio–Rad) using the Trans-Blot Turbo system (Bio–Rad) at 25V for 30 minutes. Membranes were blocked with 5% milk in Tris–Buffered Saline containing 0.05% Tween 20 (TBS–T) for 1 hour at room temperature or overnight at 4°C. Blocked membranes were incubated with primary antibodies diluted in TBS–T for 2 hours at room temperature or overnight at 4°C (Anti-KIF1A antibody, Ab271047, 1:500; Anti-VAMP2, CST-13508T, 1:1000, Anti-β–actin, A2228, 1:10000; Anti-β–tubulin, sc-80016, 1:10000). After washing, the membranes were incubated with fluorescently labeled secondary antibodies (LiCor) for 1 hour at room temperature before visualization.

### Mouse primary cell culture

Primary neuronal cell cultures were established from the brains of WT and *Sppl2b*–KO mice at embryonic day 17 (E17). Embryonic brain tissues were dissected in ice–cold Hank’s Balanced Salt Solution (HBSS, ThermoFisher, #14025092). The tissues were then incubated in HBSS supplemented with Accutase (1 ml per brain), centrifuged, and resuspended in Neurobasal medium (Gibco, #21103049) enriched with 2% B–27 (Gibco, #17504044) and 1% Glutamax (Gibco, #35050061). The suspension was filtered and plated onto wells pre–coated with Poly–D–lysine (Sigma, P6407). After 14 days in culture, neurons were fixed with 4% formaldehyde for immunofluorescence staining.

### Immunofluorescence analysis

Paraffin–embedded brain tissues were sectioned into 5 μm thick slices and mounted onto glass slides. The sections were deparaffinized by sequential washes in xylene and ethanol solutions of decreasing concentrations (99%–70%). For antigen retrieval, the slides were pressure–boiled in citrate buffer (0.1 M citric acid and 0.1 M sodium citrate) at 110°C for 5 minutes, followed by rinsing in tap water and PBS–Tween 0.05% for 5 minutes each. The sections were blocked with TNB blocking buffer (0.1 M Tris–HCl pH 7.5, 0.15 M NaCl, and 0.5% Blocking Reagent; PerkinElmer, USA) or normal goat serum (NGS; Vector Laboratories, USA) for 30 minutes at room temperature. After washing three times in PBS–T with gentle agitation, the sections were incubated overnight at 4°C with primary antibodies (Anti-KIF1A antibody, Ab271047, 1:50; Anti-VAMP2, CST-13508T, 1:1000). The following day, the sections were washed three times with PBS–T and incubated for 2 hours at room temperature with anti–mouse or anti–rabbit secondary antibodies (Alexa Fluor 488, Anti-rabbit IgG, 1:1000; Alexa Fluor 546 Anti-rabbit IgG, 1:1000). After washing, the sections were counterstained with Hoechst solution (1:500 in PBS–T) for 15 minutes with gentle agitation. The final washes were performed in PBS–T before mounting the slides with PermaFluor Aqueous Mounting Medium (ThermoScientific, USA) and allowing them to dry overnight. The prepared sections were visualized under a Nikon Eclipse E800 confocal microscope, with imaging performed using a Nikon DS–Qi2 camera at 2x, 10x, and 20x magnifications. Immunoreactive signals were quantified using ImageJ software (National Institutes of Health, MD). The fluorescence intensity of the cells was quantified using ImageJ software (National Institutes of Health, MD). Total cellular fluorescence (TCF) was calculated using the formula: TCF = Integrated Density – (Area of selected cell × Mean fluorescence of background readings). Primary neurons cells were fixed with 4% paraformaldehyde (PFA) and rinsed three times with 1X PBS. Permeabilization was performed using 0.1% Triton X–100 in 1X PBS for 10 minutes, followed by three additional PBS washes. To block nonspecific binding, cells were incubated in 3% BSA dissolved in 1X PBS for 1 hour at room temperature, then rinsed three more times with PBS. Following blocking, the fixed cells were incubated overnight at 4°C with the primary antibody diluted in 3% BSA/1X PBS. The next day, cells were washed thoroughly and incubated with a secondary antibody (1:1000 dilution) for 1 hour at room temperature. Afterward, cells were treated with Hoechst solution (Hoechst 3342, Thermo Scientific™) at a 1:500 dilution for 15 minutes at room temperature. Finally, the cells were subjected to a final wash before being mounted on Superfrost™ Plus Adhesion Microscope Slides (Epredia) using Fluoroshield™ histology mounting medium (Sigma–Aldrich).

### RNA Isolation and Quantitative PCR

Total RNA was isolated from the hippocampus and cortex of WT and *Sppl2b*–KO mice aged 12 months using the RNeasy Mini Kit (Qiagen, Cat. No. 74104), following the manufacturer’s protocol. RNA concentration and integrity were assessed with a NanoDrop ND–1000 Spectrophotometer (Thermo Fisher Scientific). One microgram of total RNA from each sample was reverse transcribed into cDNA. The reaction mixture (20 µL total volume) was prepared using the supplied master mix and amplified with an S1000 Thermal Cycler (Bio–Rad). Quantitative real–time PCR (RT–qPCR) was performed in duplicate 10 µL reactions using TaqMan Fast Advanced Master Mix (Thermo Fisher Scientific, Waltham, MA, USA) and a StepOnePlus Real–Time PCR System (Applied Biosystems, Waltham, MA, USA). The following TaqMan primer–probe sets (Thermo Fisher Scientific) were used: *Kif1a (*Mm00492863_m1), and *β*–*Actin* (Mm02619580_g1). Gene expression levels were normalized to *β*–*Actin*, and relative mRNA expression was determined using the ΔΔCt method, where fold change was calculated as ΔΔCt = ΔCt(T2) - ΔCt(T1).

## List of abbreviations

SPPL2b: Signal peptide peptidase-like 2b
WT: Wild-type
KO: Knockout
APP: Amyloid precursor protein
Aβ: Amyloid-beta
AD: Alzheimer’s disease
PD: Parkinson’s disease
KIF1A: Kinesin family member 1A
VAMP2: Vesicle-associated membrane protein 2
VAMP1: Vesicle-associated membrane protein 1
AP: adaptor proteins
AP2S1: Adaptor-related protein complex 2 sub. sigma 1
AP2A1: Adaptor-related protein complex 2 sub. alpha 1
AP1G1: Adaptor-related protein complex 1 sub. gamma 1
EEA1: Early endosome antigen 1
ACTG: Actin gamma
MA7D2: MAP7 domain-containing protein 2
APOE: Apolipoprotein E
SORL1: Sortilin-related receptor 1
HMGB1: High mobility group box 1
HMOX2: Heme oxygenase 2
SODC: Superoxide dismutase (cytosolic)
IF4G2: Eukaryotic translation initiation factor 4 gamma 2
NCAM1: Neural cell adhesion molecule 1
NSF: N-ethylmaleimide-sensitive factor
LANCL1: LanC-like protein 1
ENTP2: Ectonucleoside triphosphate diphosphohydrolase 2
LTOR1: Late endosomal/lysosomal adaptor, MAPK and mTOR activator 1
mTORC1: Mechanistic target of rapamycin complex 1
GPCR: G protein-coupled receptor
IF: Immunofluorescence
WB: Western blot.

## Declarations

### Ethics approval and consent to participate

Mice experiments ethical approval was obtained from the Stockholm Ethics Committee (15758–2019 and 12570–2021), and efforts were made to minimize the number of animals used in the study.

### Consent for publication

Not applicable

### Availability of data and materials

The datasets used and/or analysed during the current study are available from the corresponding author on reasonable request. Furthermore, proteomic dataset are available into the PRIDE website using the following account details: Username

Password: 1NAvzGl2ruLR

### Authors’ contributions

**Simone Tambaro** and **Tiziana Cabras**: Conceptualization. **Cristina Contini, Alessandra Schirru, Greca Lai, Giorgia Zodio, Jack Badman, Bjorn R.V. Bakker,** and **Ylenia Lai**: investigation. **Cristina Contini, Giacomo Diaz** and **Tiziana Cabras**. Proteomic data analysis. **Jack Badman,** and **Bjorn R.V. Bakker:** Golgi data analysis. **Simone Tambaro** and **Per Nilsson**: Mouse model generation. **Cristina Contini**, **Tiziana Cabras** and **Simone Tambaro**: Writing the original draft. **All authors**: writing review, editing and revision:

